# *Sox9* links biliary maturation to branching morphogenesis

**DOI:** 10.1101/2024.01.15.574730

**Authors:** Hannah R Hrncir, Sergei Bombin, Brianna Goodloe, Connor B Hogan, Othmane Jadi, Adam D Gracz

## Abstract

Branching morphogenesis couples cellular differentiation with development of tissue architecture. Intrahepatic bile duct (IHBD) morphogenesis is initiated with biliary epithelial cell (BEC) specification and eventually forms a heterogeneous network of large ducts and small ductules. Here, we show that *Sox9* is required for developmental establishment of small ductules. IHBDs emerge as a webbed structure by E15.5 and undergo morphological maturation through 2 weeks of age. Developmental knockout of *Sox9* leads to decreased postnatal branching morphogenesis, manifesting as loss of ductules in adult livers. In the absence of *Sox9*, BECs fail to mature and exhibit elevated TGF-β signaling and Activin A. Activin A induces developmental gene expression and morphological defects in BEC organoids and represses ductule formation in postnatal livers. Our data demonstrate that adult IHBD morphology and BEC maturation is regulated by the *Sox9*-dependent formation of precursors to ductules during development, mediated in part by downregulation of Activin A.

## INTRODUCTION

Developmental morphogenesis establishes tissue architecture and ensures proper function into adulthood. The mammalian liver is highly organized and supports anatomical compartmentalization of different physiological and biochemical functions ^1^. Bipotential epithelial progenitor cells, termed hepatoblasts, give rise to hepatocytes and biliary epithelial cells (BECs) during liver development ^2^. Early BECs differentiate in response to TGF-β, Notch, and Yap/Hippo signaling while hepatocyte specification is believed to be a default fate of hepatoblasts in the absence of pro-BEC signaling ^3–8^. BECs line the intrahepatic bile ducts (IHBDs), which are hierarchical in structure. Small, peripheral “ductules” drain into large “ducts” that eventually join the extrahepatic bile duct ^9^.

Mechanisms linking BEC specification and *de novo* bile duct formation have been well-studied, but less is known about the contribution of pre-existing BECs to IHBD expansion. IHBD morphogenesis progresses over time from the liver hilum towards the periphery, with BEC markers first detectable between E11.5-E13.5 ^10,11^. As BECs specify, they organize around the portal vein to form a ductal plate ^2^. Conventional models propose that ductal plates, which are “stacked” on top of one another, give rise to discontinuous primitive ductal structures that interconnect and elongate through a process termed tubulogenesis ^2,12^. These structures rearrange and regress through late embryonic and early postnatal development to give rise to adult IHBDs ^12–14^. However, the genetic mechanisms underlying developmental specification of duct-ductule hierarchy and global IHBD architecture remain poorly understood.

Branching morphogenesis, which includes tubulogenesis, has been the predominant framework for understanding how ramified epithelium expands into surrounding tissue. It can be broadly classified as: (i) stereotyped, where highly conserved patterns of epithelial branching correlate with overall tissue growth, or (ii) stochastic, which is initially sporadic and dependent on epithelial remodeling to establish homeostasis ^15^. Stochastic branching is observed in tissues where ductal structures must “invade” a pre-existing parenchyma, like the pancreas ^16^. Since developing IHBDs migrate through the hepatic parenchyma, principles of stochastic branching morphogenesis may also govern their formation ^17,18^.

The BEC-associated transcription factor SOX9 is a key regulator of stereotyped and stochastic branching in the lung, salivary gland, and thyroid ^19–22^. SOX9 is activated in the ductal plate at E11.5 in response to pro-BEC signals and remains expressed in all BECs throughout adulthood ^10,23^. Surprisingly, conditional deletion of *Sox9* in hepatoblasts (Sox9cKO) has been reported to have minimal impact on IHBD development, resulting only in delayed BEC specification from E15.5-P6 ^10,24^. However, subsequent studies of adult Sox9cKO mice identified defects in BEC polarity and loss of primary cilia, with aged mice developing liver cysts ^25^. These data raise compelling questions about the importance of *Sox9* in regulating BEC identity and whether it may contribute to IHBD branching morphogenesis.

Using quantitative, whole-tissue 3D imaging of IHBDs, we find that loss of *Sox9* during development disrupts branching morphogenesis, leading to reduced numbers of ductules in adult livers. Transcriptomic assays demonstrate that Sox9cKO BECs resemble developing BECs and continue to express high levels of NCAM1, which is normally downregulated after development. Early postnatal livers exhibit decreased IHBD complexity in the absence of *Sox9*, demonstrating that adult ductule defects originate during development. 3D imaging of normal IHBD development suggests that ducts do not form from progressive merging of disconnected cysts, but rather through advancing growth and remodeling of an interconnected biliary “web” that is dependent on *Sox9*. Mechanistically, *Sox9* contributes to IHBD branching morphogenesis, in part by suppressing Activin A. Together our studies identify two major components of IHBD morphogenesis: (1) *Sox9*-independent morphogenesis of large ducts and (2) Activin A-inhibited, *Sox9* driven morphogenesis of small ductules.

## RESULTS

### Ductal paucity in adult Sox9cKO mice

To investigate how developmental loss of *Sox9* impacts IHBD biology, we crossed *Sox9*-floxed mice to *Albumin*-Cre mice to generate mice with conditional deletion of *Sox9* in hepatoblasts (*Sox9^fl/fl^:Alb^Cre^*; Sox9cKO). Sox9cKO mice undergo Cre recombination at E10.5, resulting in loss of *Sox9* in hepatocytes and BECs ^10,26^. To avoid confounding effects of rare hepatic cystogenesis, which was previously reported in Sox9cKO mice as early as 24 weeks of age, we only studied adult mice between 6-12 weeks ^25^. We confirmed loss of SOX9 in adult Sox9cKO livers by co-localizing SOX9 with the BEC-specific marker EpCAM by immunofluorescence (IF) (Figure 1A). Previous reports have described loss of primary cilia in adult and developing BECs following conditional ablation of *Sox9* using both *Alb^Cre^* and *Alfp^Cre^* alleles ^24,25^. We confirmed loss of primary cilia in our adult Sox9cKO mice by IF for acetylated α-tubulin (Ac-tubulin) (Figure S1A). In contrast with previous reports that loss of *Sox9* only delays BEC specification, we observed reduced numbers of both EpCAM+ BECs and distinct ductal structures per portal field in Sox9cKO mice (Figure 1B) ^10,24^. To determine if ductal paucity results from decreased BEC proliferation, we performed co-IF for EpCAM and Ki-67. Proliferation rates were unchanged between control and Sox9cKO BECs (Figure S1B). Since BECs are largely quiescent during homeostasis, we further assayed proliferative potential by inducing liver injury with thioacetamide (TAA) for 1 month, which results in extensive proliferation of pre-existing BECs ^27^. Once again, we saw no change in BEC proliferation in Sox9cKO mice (Figure S1C). To determine if apoptosis contributes to ductal paucity in Sox9cKO mice, we conducted co-IF for EpCAM and cleaved caspase-3. We did not detect apoptosis during homeostasis in either control or Sox9cKO BECs (n = 3 biological replicates per control and Sox9cKO, 10 portal fields quantified per replicate; data not shown). Therefore, adult Sox9cKO mice exhibit ductal paucity that is not a result of decreased proliferation or increased apoptosis.

**Figure 1.**
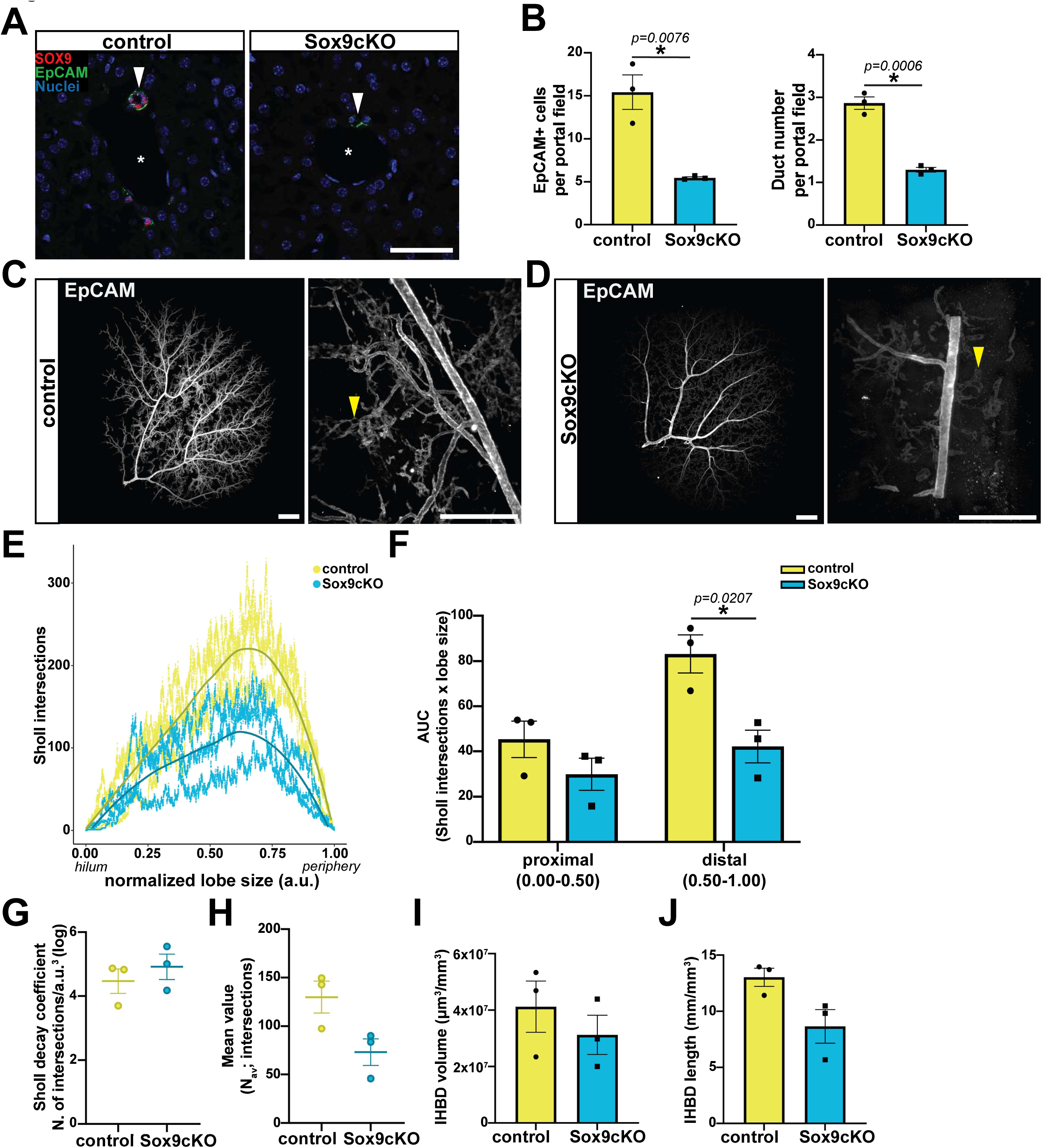
Loss of Sox9 during development results in ductal paucity in adult mice. **(A)** IF of adult control and Sox9cKO mice confirms loss of *Sox9* in Sox9cKO mice compared to control (scale bar represents 50μm; white asterisks indicate portal vein). **(B)** Sox9cKO mice exhibit a loss of BECs and reduced IHBDs per portal field (n = 3 biological replicates per group). **(C)** Whole tissue imaging of control liver lobes illustrates complex IHBD hierarchy, while **(D)** Sox9cKO mice exhibit significant loss of ductules, IHBD discontinuity, and reduced EpCAM expression (scale bars represent 0.5mm; arrowheads mark representative ductules). **(E)** Sholl analysis demonstrates quantitative loss of branching complexity throughout Sox9cKO IHBDs (arbitrary units, a.u., represent distance normalized to lobe size). **(F)** AUC of proximal (0.00-0.50a.u.) vs. distal (0.50-1.00a.u.) Sholl intersections reveals significant loss of branching in distal IHBDs. **(G)** Sholl decay coefficient is unchanged between control and Sox9cKO IHBDs and **(H)** the mean value of total Sholl intersections (N_av_) trends downward (*p* = 0.056). Total **(I)** volume and **(J)** length are not significantly different between control and Sox9cKO IHBDs (n = 3 left lobes per group).

### *Sox9* is required for proper IHBD morphology

To determine if ducts or ductules are differentially sensitive to the loss of *Sox9*, we performed whole-tissue imaging using iDISCO+ tissue clearing and light sheet microscopy of adult control and Sox9cKO livers ^28^. Control tissues clearly demonstrated expected IHBD hierarchy, with large ducts at the hilum decreasing in size towards the periphery and terminating in web-like ductules (Figure 1C, Video S1 & S2). In contrast, Sox9cKO IHBDs were disorganized and discontinuous, with a general loss of peripheral webbing (Figure 1D, Video S1 & S2). Ductules in Sox9cKO tissues also stained much more dimly for EpCAM than controls (Figure 1D, Video S2; arrowheads).

To gain a better understanding of Sox9cKO ductal paucity and overall IHBD morphology, we performed quantitative image analysis of 3D IHBD imaging data. We reasoned that Sholl analysis, which is routinely applied to describe neuron branch complexity, could be applied to quantify global IHBD morphology ^29^. 3D Sholl analysis of IHBDs was carried out by defining the hilum as the starting point, quantifying branch point intersections at three-dimensional 1µm spherical intervals, and normalizing distance from the hilum to total liver size. IHBD branching, as represented by Sholl intersections, was reduced in Sox9cKO livers, consistent with qualitative observations of small ductule loss (Figure 1E). To determine if ductal paucity was localized to distinct IHBD regions, we performed area under the curve (AUC) analysis on Sholl intersections. We found that proximal (0.00-0.50 arbitrary units; a.u.) branching complexity was minimally impacted, while distal AUC (0.50-1.00a.u.) was significantly decreased in Sox9cKO livers, consistent with the loss of peripheral ductules (Figure 1F). To further quantify *Sox9* regulated changes in IHBD architecture, we carried out regression and curve fitting analyses to characterize branching properties across samples, as previously described for analysis of neurons ^30,31^. Sholl decay coefficients, which describe the rate of change in branch density across a structure, were calculated from linear regression of semi-log plots of intersections vs. normalized distance (Figure S1D) ^32^. Consistent with AUC, Sholl decay was increased in Sox9cKO samples, reflecting decreased branch density in the liver periphery (Figure 1G, Figure S1D). Ductal paucity in adult mice was further supported by a downward trend in the average number of Sholl intersections (N_av_) in Sox9cKO mice (Figure 1H). We also compared the maximum number of Sholl intersections and the critical value (N_m_), which is the maximum of the polynomial function and therefore accounts for technical noise in biological samples, for control and Sox9cKO livers ^30,31^. We found a significant decrease in both values in Sox9cKO mice, demonstrating a decrease in maximum branch point intersections (Figure S1E). To determine the distance at which maximum branching occurs, we analyzed the critical radius (r_c_). Loss of peripheral branching in Sox9cKO mice is reflected by a slight shift in r_c_ toward the hilum (Figure S1F). However, this difference does not reach statistical significance, which may suggest limitations in global Sholl measurements to identify localized phenotypes. To quantify overall IHBD size independent of branching morphology, we measured total IHBD volume and length, and normalized these values to overall liver lobe volume. While the total length of Sox9cKO IHBDs trended downward, overall volume was unchanged (Figure 1I, & J). This may be due to the relative contribution of large ducts, which are present and dilated in Sox9cKO mice, to total IHBD volume and length. Collectively, quantitative whole tissue imaging reveals that ductal paucity in adult Sox9cKO mice is primarily characterized by reduced numbers of peripheral ductules.

### Sox9cKO BECs exhibit elevated TGF-β signaling and transcriptomic features of immaturity

To determine mechanisms linking loss of *Sox9* to ductal paucity, we performed bulk RNA-seq of FACS-isolated BECs from adult control and Sox9cKO mice (Figure S2A). As expected, control and Sox9cKO samples clustered by genotype on PCA (Figure S2B). We identified 766 upregulated and 422 downregulated genes in Sox9cKO BECs (q-value < 0.05; LFC > 1) (Figure 2A & Table S1). Several of the top upregulated genes in Sox9cKO BECs are known to be involved in TGF-β signaling, consistent with prior reports of increased TβRII expression in developing BECs of *Sox9-*deficient livers (Figure S2C) ^10^. To determine if TGF-β is broadly upregulated, we conducted gene set enrichment analysis (GSEA), which demonstrated significant enrichment of the TGF-β signaling pathway in Sox9cKO BECs (Figure 2B). To validate elevated TGF-β signaling, we performed IF for pSMAD2, which was elevated in Sox9cKO BECs (Figure 2C). We next examined the expression of all known TGF-β superfamily ligands, to ask if a potential autocrine mechanism could explain increased pSMAD2 in Sox9cKO BECs (Figure S2D). Only *Inhba*, the gene precursor to Activin A, was elevated in Sox9cKO mice by RNA-seq. We confirmed this result by IF, which demonstrated substantially increased Activin A in EpCAM+ BECs of Sox9cKO livers (Figure 2D). Interestingly, Activin A is known to inhibit branching morphogenesis in salivary glands, kidney, and prostate epithelium *in vitro* ^33,34^. Previous studies have also suggested that Activin A plays an important role in promoting BEC specification, although its precise role remains unclear ^3^. To examine the cell-type specific expression of *Inhba* coincident with BEC specification, we leveraged recently published transcriptomic data from developing hepatoblasts, hepatocytes, and BECs ^35^. We confirmed that *Inhba* is expressed at low levels in hepatoblasts and increases over time in both embryonic hepatocytes and embryonic BECs, supporting a potential role for Activin A in hepatoblast differentiation (Figure S2E).

**Figure 2.**
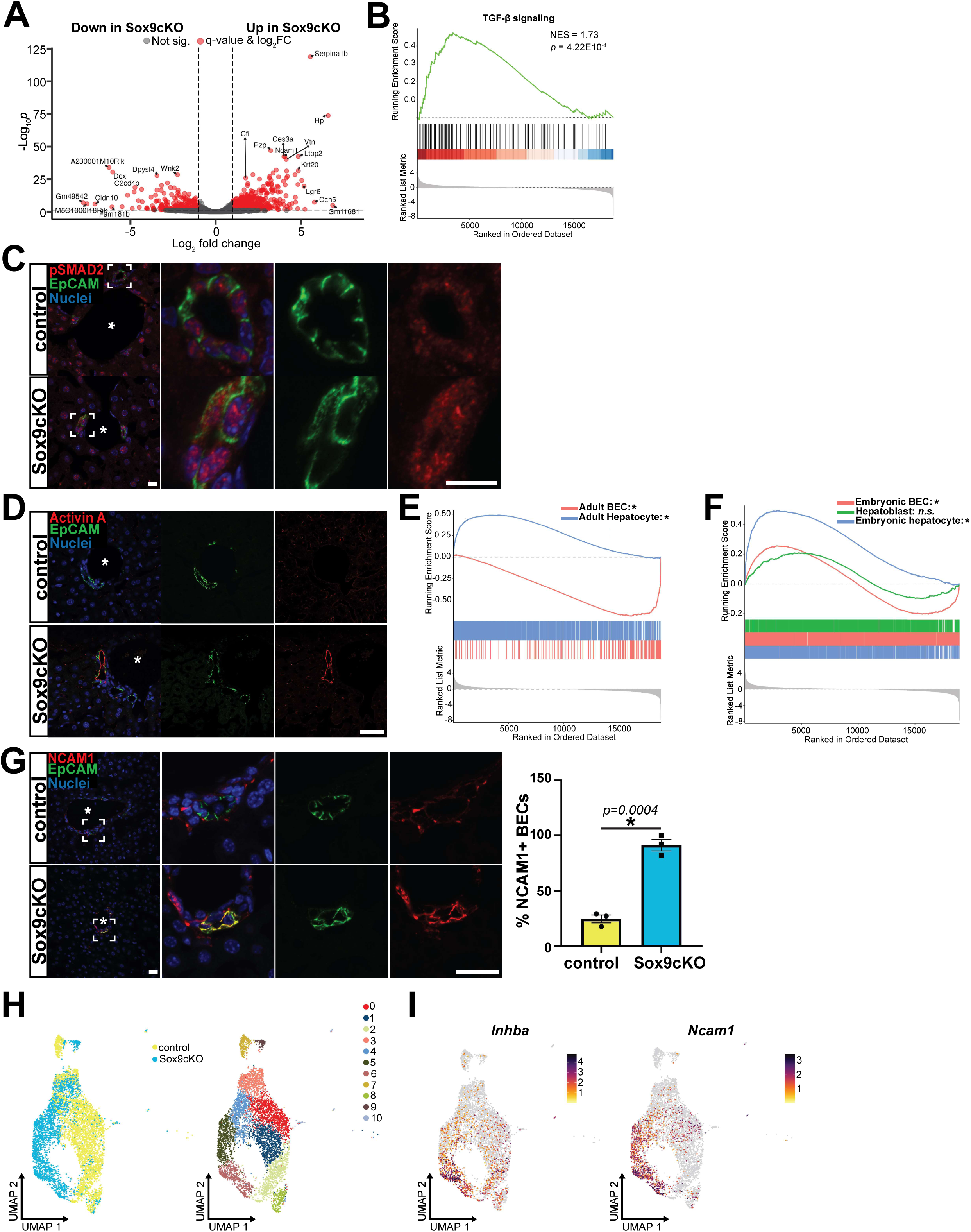
Sox9 promotes biliary identity and limits TGF-β signaling in BECs. **(A)** Bulk RNA-seq identifies DEGs between Sox9cKO and control BECs. **(B)** GSEA demonstrates significant enrichment of TGF-β signaling in Sox9cKO BECs relative to control BECs. **(C)** Co-localization of pSMAD2 and EpCAM confirms elevated TGF-β signaling in Sox9cKO BECs (scale bar represents 5μm; white asterisks indicate portal vein). **(D)** IF demonstrates elevated Activin A in EpCAM+ BECs in Sox9cKO livers (scale bar represents 50μm; white asterisks indicate portal vein). **(E)** GSEA demonstrates that Sox9cKO BECs are de-enriched for adult BEC gene signature (NES: −2.70; *p* = 3.67E-10) and enriched for adult hepatocyte gene signature (NES: 2.31; *p* = 3.67E-10). **(F)** GSEA of liver development gene sets shows that the Sox9cKO BEC transcriptome resembles embryonic BECs (NES: 1.23; *p* = 6.06E-6) and embryonic hepatocytes (NES: 2.32; *p* = 3.00E-10), but not hepatoblasts (NES: 0.99; *p =* 0.55). **(G)** Co-localization of NCAM1 and EpCAM is significantly elevated in Sox9cKO compared to controls (scale bar represents 5μm; white asterisks indicate portal vein). **(H)** scRNA-seq of FACS-isolated EpCAM+ BECs reveals subpopulation-specific transcriptomic changes in Sox9cKO. **(I)** Upregulation of *Inhba* and *Ncam1* in Sox9cKO samples is specific to Sox9cKO ductule-like cells (cluster 6).

Since TGF-β signaling participates in early BEC specification, we next asked if Sox9cKO-associated ductal paucity was associated with long-term defects in BEC specification or maturation. GSEA of published gene signatures from adult mice demonstrated that Sox9cKO BECs were enriched for hepatocyte-associated genes and de-enriched for BEC associated genes, suggesting a loss of BEC identity (Figure 2E) ^36^. We reasoned that loss of *Sox9* at E10.5 in the Sox9cKO model could result in incomplete BEC maturation or an inability to suppress the “default” hepatocyte differentiation program associated with liver development ^8,11^ To further determine if Sox9cKO BECs are more embryonic-like, we performed GSEA with published RNA-seq from: (1) E12.5-E14.5 hepatoblasts, (2) E15.5-E17.5 BECs, and (3) E15.5-E17.5 hepatocytes ^35^. Embryonic BEC and embryonic hepatocyte gene signatures were significantly enriched in adult Sox9cKO BECs, but hepatoblast genes were not, suggesting that *Sox9* is required for maturation of the BEC transcriptome, but dispensable for loss of hepatoblast identity (Figure 2F). To validate incomplete BEC maturation, we performed IF for NCAM1, which has been reported to be expressed exclusively by developing or injured BECs and was highly elevated in Sox9cKO BECs by RNA-seq (Figure S2C) ^37^. While co-localization of NCAM1 and EpCAM demonstrated that 24.5 ± 3.5% of BECs in healthy, adult controls are NCAM1+, this number rose to 91.4 ± 5.2 % in Sox9cKO mice (Figure 2G). Taken together, these data suggest that Sox9cKO BECs are transcriptomically immature and have gene expression signatures consistent with a failure to fully acquire BEC identity. Additionally, these defects may be driven by upregulation of Activin A, a ligand shown in other tissues to repress branching and a proposed regulator of early BEC specification.

### *Sox9* has subpopulation-specific impacts on BEC gene expression

Based on the loss of ductules in Sox9cKO livers by whole-tissue imaging, we hypothesized that defects observed in Sox9cKO BECs could be subpopulation-specific. To this end, we performed scRNA-seq on FACS-isolated EpCAM+ BECs from control and Sox9cKO mice. We identified 11 distinct clusters across 6,152 BECs (3,498 control; 2,654 Sox9cKO) (Figure 2H, S3A, & Table S2). Most control and Sox9cKO BECs clustered separately, with the exception of clusters 3, 8, and 10, which contained a mixture of control and Sox9cKO BECs (Figure 2H, S3B). Next, we wanted to determine if one or more subpopulations of BECs identified by scRNA-seq correlated with large ducts or ductules. We started by examining gene expression of previously reported biomarkers of ducts (*Nfatc3, Slc10a2, Cftr, Slc4a2, Sctr,* and *Mtnr1a*) and ductules (*Hrh1, Sp1, Nfatc4,* and *Nfatc2*), in our scRNA-seq data ^38–43^. These markers were either non-specific and expressed broadly across all clusters, or not detected in a sufficient number of cells to infer duct/ductule identity (Figure S3C). Recent studies from our lab utilized a Sox9^EGFP^ reporter transgene to characterize BEC heterogeneity, finding that IHBDs contain subpopulations of Sox9^EGFP-low^ and Sox9^EGFP-high^ cells. Sox9^EGFP-high^ cells are more prevalent in ductules, suggesting that BECs with similar gene signatures are also more likely to be located in ductules ^23^. GSEA demonstrated enrichment of Sox9^EGFP-high^ gene signature in clusters 1, 2, 5, 6, and 8, suggesting ductule identity (Figure S3D). Therefore, we classified clusters 1, 2, 5, 6, and 8 as belonging to ductules and clusters 0, 3, 4, 7, 9 & 10 as belonging to ducts (Figure S3E). To assess whether ducts or ductules are more impacted by loss of *Sox9*, we performed differential gene expression analysis between control and Sox9cKO samples, within ductule and duct groups. We found 1,860 differentially expressed genes (DEGs) between control and Sox9cKO duct-like BECs and 1,830 DEGs between control and Sox9cKO ductule-like BECs (Figure S3F). Interestingly, gene expression analysis also identified *Ncam1* and *Inhba* upregulation specifically in the Sox9cKO ductule cluster (Figure 2I). Taken together, scRNA-seq suggests that loss of *Sox9* has transcriptomic impacts on duct and ductule BECs, but that increased *Inhba* and *Ncam1* upregulation observed by bulk RNA-sequencing is likely derived from a ductule-resident subpopulation of Sox9cKO BECs. These data support our observation of a ductule phenotype in Sox9cKO mice by whole-tissue IF, and further implicate elevated *Inhba*/Activin A in Sox9cKO-associated phenotypes.

### *Sox9* is required for normal biliary organoid formation and cystic morphology

We next wanted to determine if *Sox9* is also required for the establishment of biliary organoids (referred to from here forward as “mouse intrahepatic cholangiocyte organoids” or “mICOs” for consistency with established nomenclature) ^44^. We reasoned that mICOs would also provide a highly tractable platform for dissecting mechanisms related to the phenotypic impact of *Sox9* in BECs. FACS-isolated control and Sox9cKO BECs were plated as single cells and allowed to grow in standard mICO media conditions for 7d prior to analysis ^45^. Compared to controls, Sox9cKO BECs formed approximately 50% fewer mICOs (Figure 3A & B). More strikingly, a majority of Sox9cKO mICOs failed to form the clearly defined lumens observed in control samples and characteristic of ductal epithelial organoids (Figure 3A & C) ^45^. We confirmed that these mICOs, which we termed “non-cystic”, continue to be viable by conducting live cell imaging (Video S3). Despite non-cystic morphology, we observed no change in area between control and Sox9cKO mICOs at 7d of culture, consistent with our observation that proliferation is unaffected by loss of *Sox9 in vivo* (Figure 3D). To directly assay mICO proliferation, we conducted and quantified whole mount IF for Ki-67 and saw no change in the percent of proliferating BEC nuclei between control and Sox9cKO mICOs (Figure S4A). Together, these results establish that *Sox9* is required for mICO formation and organization from single BECs *in vitro*.

**Figure 3.**
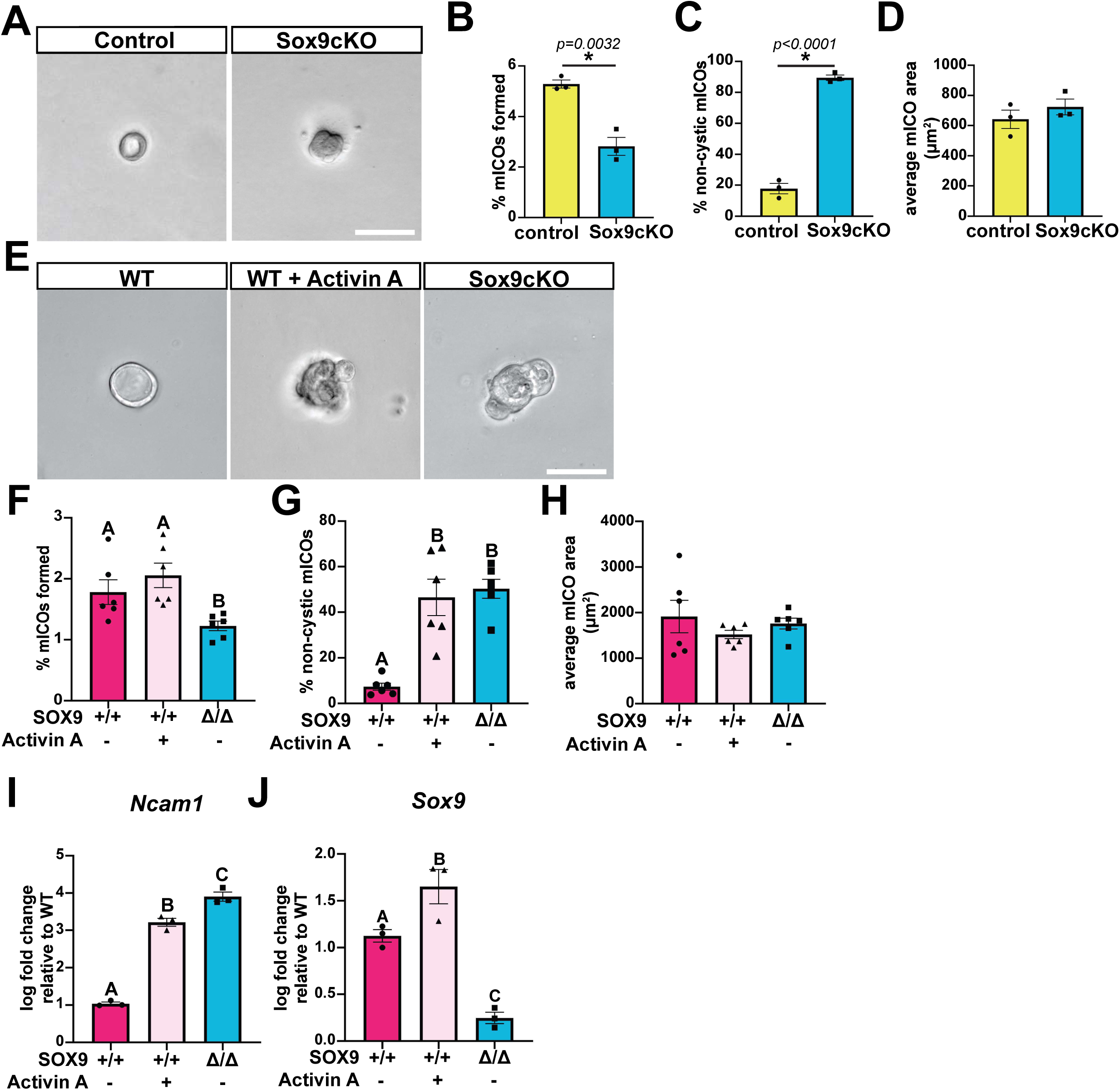
Sox9 promotes cystic morphology in BEC-derived organoids. **(A)** Single Sox9cKO BECs cultured for 7d fail to form lumens typical of control mICOs. **(B)** Sox9cKO BECs form significantly fewer mICOs, a majority of which **(C)** exhibit non-cystic morphology. **(D)** mICO area is unchanged between control and Sox9cKO. **(E)** 50ng/mL Activin A is sufficient to induce Sox9cKO morphology in WT BECs, but **(F)** unlike Sox9cKO (Sox9 Δ/Δ) did not impact mICO organoid formation. **(G)** Rates of non-cystic mICO morphology were similar between Sox9cKO and Activin A treated WT BECs. **(H)** Organoid area remained unchanged between all groups. **(I)** *Ncam1* was significantly upregulated in Sox9cKO mICOs by RT-qPCR, and induced by Activin A in WT mICOs. **(J)** Activin A upregulates *Sox9* expression in WT mICOs (letters indicate grouping by significance*, p* < 0.05; scale bars represent 50μm).

### Activin A signaling regulates non-cystic mICO phenotype

Next, we sought to determine if elevated Activin A identified *in vivo* was responsible for the non-cystic phenotype observed in Sox9cKO mICOs. We isolated wild-type (WT) BECs from C57Bl/6 mice and plated them in standard mICO conditions with or without Activin A for 7d prior to analysis. Sox9cKO BECs were isolated in parallel as a positive control for non-cystic morphology. Treating WT BECs with Activin A did not recapitulate loss of organoid formation seen in Sox9cKO mICOs (Figure 3E & F). However, Activin A treated WT mICOs exhibited non-cystic morphology at approximately the same rate observed in Sox9cKO mICOs, establishing that exogenous Activin A is sufficient to interfere with normal mICO morphology (Figure 3E & G). Activin A did not impact mICO size (Figure 3H). To determine if Activin A is also sufficient to induce immature BEC gene expression *in vitro*, we performed RT-qPCR. As observed in Sox9cKO BECs *in vivo*, *Ncam1* was significantly upregulated by Activin A in mICOs (Figure 4I). Interestingly, we also found that *Sox9* was upregulated in Activin A treated mICOs (Figure 4J). These data demonstrate that exogenous Activin A partially recapitulates Sox9cKO phenotypes in WT mICOs, inducing non-cystic morphology but failing to decrease organoid formation rates. Activin A also upregulates *Ncam1* and *Sox9* in mICOs, suggesting a link between Activin A and gene regulatory defects observed in Sox9cKO IHBDs *in vivo*.

**Figure 4.**
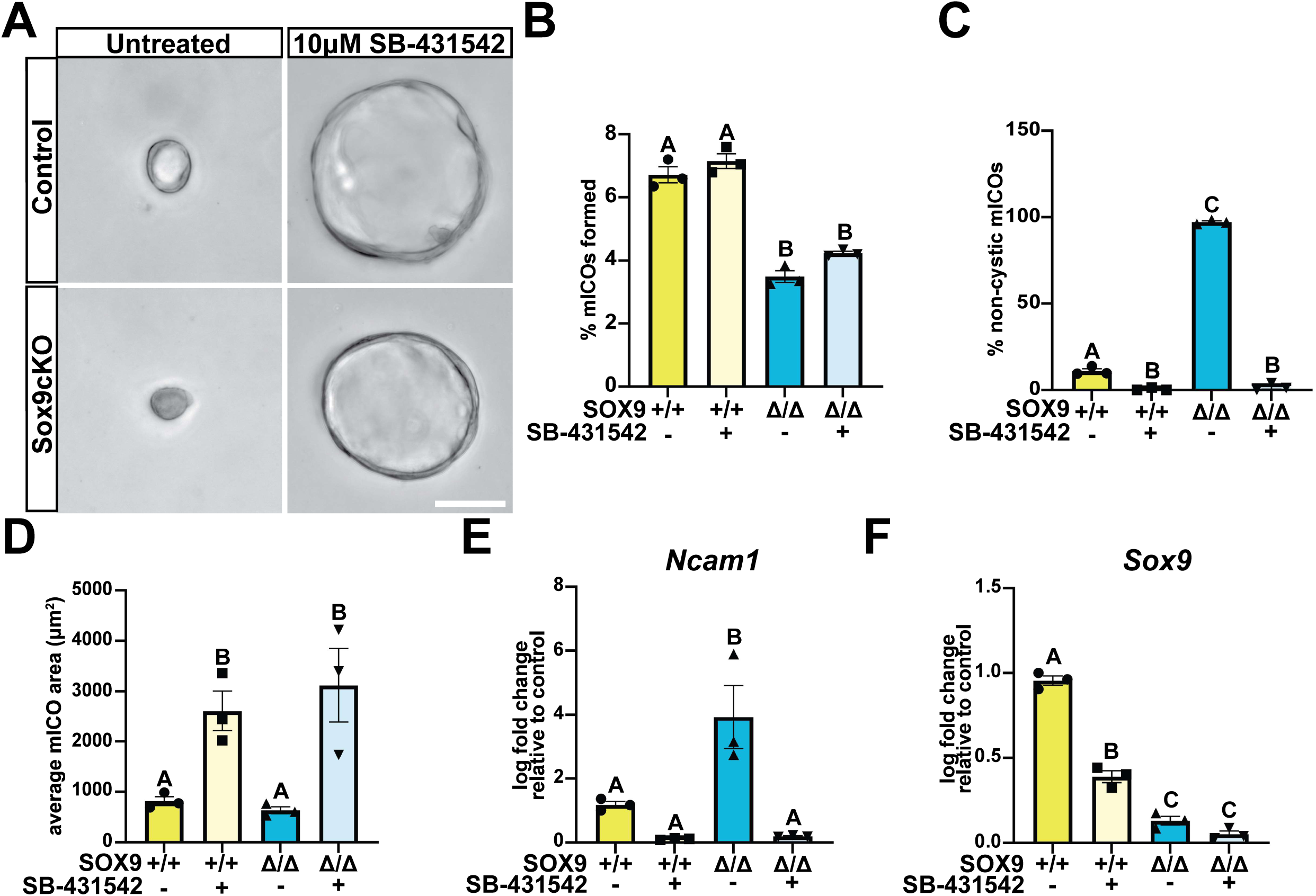
Inhibiting Activin A signaling rescues biliary organoid morphology. **(A)** Culturing single BECs with 10μM SB-431542 for 7d rescues non-cystic morphology in Sox9cKO mICOs (scale bar represents 50μm). **(B)** Decreased mICO formation by Sox9cKO BECs is not impacted by SB-431542, but **(C)** non-cystic morphology is completely rescued. Notably, rare non-cystic mICOs in the control group are also significantly reduced. **(D)** SB-431542 treatment increases mICO size independent of *Sox9*. **(E)** RT-qPCR demonstrates that SB-431542 downregulates *Ncam1* in control and Sox9cKO mICOs and **(F)** *Sox9* in control mICOs (letters indicate grouping by significance, *p* < 0.05).

To confirm that canonical downstream Activin A signaling is responsible for non-cystic mICO morphology, we attempted a functional rescue by inhibiting signaling from activin type II receptors. SB-431542 is a widely used, specific inhibitor of ALK4/5/7 that prevents phosphorylation of SMAD2/3 ^46^. We isolated and cultured single control and Sox9cKO BECs for 7d in standard mICO conditions with or without SB-431542. ALK4/5/7 inhibition did not impact the number of mICOs formed (Figure 4A & B). Notably, non-cystic mICO morphology in Sox9cKO samples was completely rescued by SB-431542 (Figure 4A & C). SB-431542 also increased organoid size comparably in both control and Sox9cKO mICOs (Figure 4A & D). Next, we conducted RT-qPCR to determine whether ALK4/5/7 inhibition had the opposite effect on *Ncam1* and *Sox9* expression as Activin A treatment. *Ncam1*, which was elevated in untreated Sox9cKO BEC organoids relative to controls, was significantly downregulated in both control and Sox9cKO mICOs following SB-431542 treatment (Figure 4E). Additionally, *Sox9* was significantly downregulated in control organoids following SB-431542 treatment, reinforcing the link between Activin A and induction of *Sox9* (Figure 4F). These results demonstrate that excess ALK4/5/7-mediated signaling is responsible for the non-cystic morphology observed in Sox9cKO mICOs. Additionally, signaling through ALK4/5/7 positively regulates *Sox9* and *Ncam1* expression. Together with *in vivo* data demonstrating increased *Inhba* and Activin A expression in Sox9cKO BECs, our mICO data suggest that Activin A signaling drives a negative feedback loop by upregulating *Sox9*.

### Late IHBD development proceeds via proliferative expansion of a continuous ductal “web”

To identify timepoints relevant to the role of Activin A and *Sox9* in establishing ductule networks, we next applied whole-tissue imaging to IHDB development. We reasoned that the ability to detect early BECs at high sensitivity and with minimal manipulation of the tissue using iDISCO+ and light sheet microscopy could clarify key events in IHBD development. We imaged EpCAM+ ductal structures in wild-type, C57Bl/6 livers at E15.5, E17.5, P1, P5, and P14. Consistent with previous reports, EpCAM+ structures formed at the hilum at early developmental timepoints and progressively advanced toward the periphery through P5 (Figure 5A) ^12^. Surprisingly, we noted very little discontinuity between BECs across IHBD development, which was previously reported through E17 ^12^. While rare peripheral ductal structures were disconnected from the main mass of EpCAM+ cells at E15.5, no disconnected structures were observed at E17.5 or later (Figure 5A & B). In contrast to reports relying on retrograde ink injection, ductal structures were present as a tight “web” lacking a clear hierarchy of ducts and ductules from E15.5 through P1 (Figure 5A & B) ^12^. While well-defined duct-ductule hierarchy was still lacking at P1, a small subset of ducts began to exhibit slightly enlarged size while remaining tightly intertwined in the IHBD web (Figure 5C). By P5, large ducts webbed with smaller, peripheral ductule branches can be observed extending away from the hilum towards the periphery of the liver (Figure 5A). Surprisingly, even by P14, a majority of IHBD branches continue to be webbed and lack the distinct, interlobar large ducts that are characteristic of adult IHBDs (Figure 1C & 5A). At P14, minimally webbed large ducts appear to be emergent at the hilum and expanding towards the periphery, suggesting a progressive remodeling of IHBD hierarchy over time (Figure 5D). These data further establish that IHBD morphogenesis is a dynamic postnatal process, with IHBDs starting to demonstrate a clear hierarchy by P5, but not resembling a fully mature interlobar network even by 2 weeks of age.

**Figure 5.**
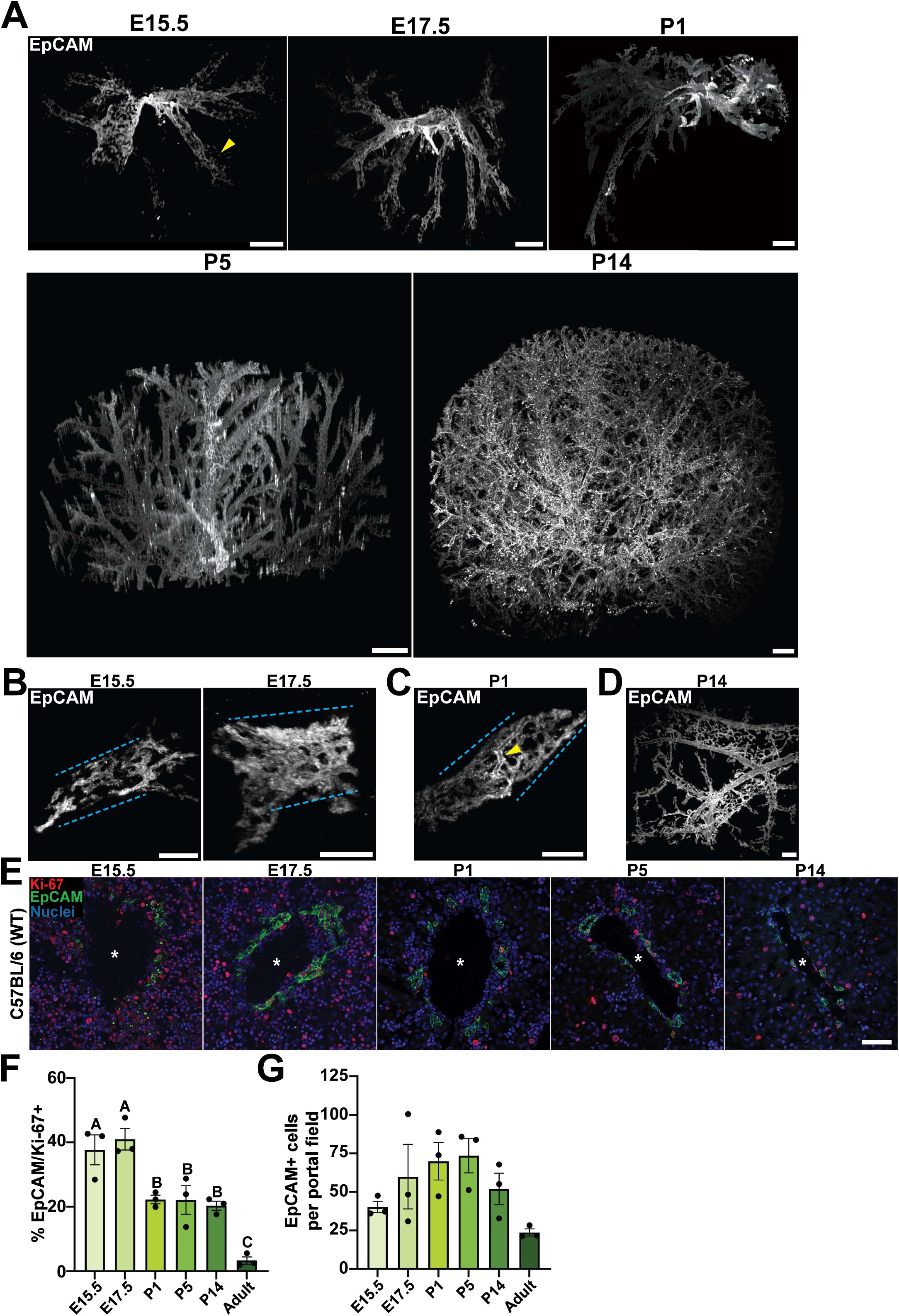
Whole tissue imaging of EpCAM+ IHBD development. **(A)** IHBDs develop primarily as an interconnected web of EpCAM+ BECs from E15.5 through P14, originating at the hilum and extending to the periphery by P1, with rare disconnected BECs visible closer to the periphery at E15.5 (yellow arrowhead marks non-continuous BECs; scale bar represents 0.5mm). **(B)** High-magnification images demonstrate discontinuous BECs at E15.5, which are absent by E17.5 **(C)** At P1, early duct-ductule hierarchy begins to emerge as some IHBDs become enlarged but remain intertwined with web-like peripheral IHBDs. **(D)** Duct-ductule hierarchy is clearly established at P14, but large hilar ducts continue to be surrounded by web-like peripheral IHBDs (scale bars represent 150μm; yellow arrowhead denotes enlarged IHBD; dashed blue lines denote portal veins). **(E)** Co-localization of Ki-67 and EpCAM reveals proliferative BECs throughout IHBD development (scale bar represents 50μm; white asterisks indicate portal vein). **(F)** Quantification of BEC proliferation reveals abundant embryonic proliferation and decreasing, but high proliferation rates relative to adult tissue during postnatal IHBD development. **(G)** EpCAM+ BECs per portal field trend upwards E15.5-P5 until decreasing P14 and into adulthood.

Models for IHBD development focus on BEC specification from hepatoblasts, but the proliferative expansion of BECs post-specification is less well-characterized. Some have proposed that IHBD tubulogenesis occurs in the absence of BEC proliferation ^17^. To determine the contribution of BECs to IHBD growth during development, we co-localized EpCAM and Ki-67 at the same timepoints assayed by whole-tissue imaging. During embryonic development, EpCAM+ BECs are hyperproliferative, with 37.7 ± 4.6% of BECs expressing Ki-67 at E15.5 and 41.0 ± 3.4% at E17.5 (Figure 5E & F). Postnatally, BEC proliferation drops to ∼20% at P1 through P14 (Figure 5E & F). A final decrease in BEC proliferation occurs between P14 and adulthood with 3.30 ± 1.1% of BECs proliferating at 8 weeks of age, consistent with proliferation rates observed in adult control and Sox9cKO livers (Figure S1A, 5E & F). To determine if there is a relationship between BEC proliferation and IHBD expansion, we quantified BECs per portal field using conventional IF. BEC numbers followed a similar trend to proliferation, initially increasing in number and then trending downwards postnatally (Figure 5E & G). This downward trend is coincident with and likely explained by postnatal IHBD remodeling and emergence of duct-ductule hierarchy. These data suggest that BECs are key contributors to IHBD expansion during development following hepatoblast differentiation, and that regulatory pathways controlling postnatal IHBD development may have substantial impacts on adult IHBD morphology.

### Loss of *Sox9* results in early postnatal ductal paucity

Whole-tissue imaging revealed that P5 IHBDs consist of highly proliferative, ductule-like webs, while also demonstrating the earliest signs of duct-ductule hierarchy. Therefore, to study the impact of *Sox9* on IHBD morphogenesis during development, we collected livers from P5 control and Sox9cKO mice. Using conventional IF on tissue sections, we found that P5 Sox9cKO mice exhibit ductal paucity compared to control livers, consistent with adult samples (Figure 6A & B). The percent of Ki-67+/EpCAM+ BECs was unchanged between P5 Sox9cKO and control tissues, further suggesting that ductal paucity in Sox9cKO livers is not caused by reduced proliferation, even in developing IHBDs (Figure 6A & C). To examine global IHBD morphology during development, we subjected P5 livers to iDISCO+ and light-sheet imaging. Similar to large ducts in adult mice, P5 Sox9cKO mice exhibited main duct branches extending from the hilum to the periphery, with no apparent morphological defects (Figure 6D). However, these main branches were largely devoid of the web-like peripheral IHBDs observed at P5 in WT and control livers (Figure 6D). Sholl analysis revealed a pronounced loss of IHBD complexity in P5 Sox9cKO mice, in contrast to the more subtle differences in branching observed between control and Sox9cKO adults (Figure 6E & S5A). This result was confirmed by large decreases in both proximal (0.0-0.5) and distal (0.5-1.0) AUC in Sox9cKO livers (Figure 6F). Sholl decay was unchanged between P5 control and Sox9cKO livers, suggesting a uniform branch density throughout the tissue (Figure 6G). The universal loss of branching in P5 Sox9cKO tissue was further apparent in reduced average branch density, as well as the actual and calculated maximum number of branch point intersections (Figure 6H & S5B). The distance at which maximum branching occurs (r_c_) was unchanged between control and Sox9cKO, further reinforcing global changes in IHBD morphology in developing Sox9cKO livers (Figure S5C). Together, these data demonstrate that ductal paucity in adult Sox9cKO mice originates during development and suggest that *Sox9* is critical for morphogenesis of web-like precursors to ductules.

**Figure 6.**
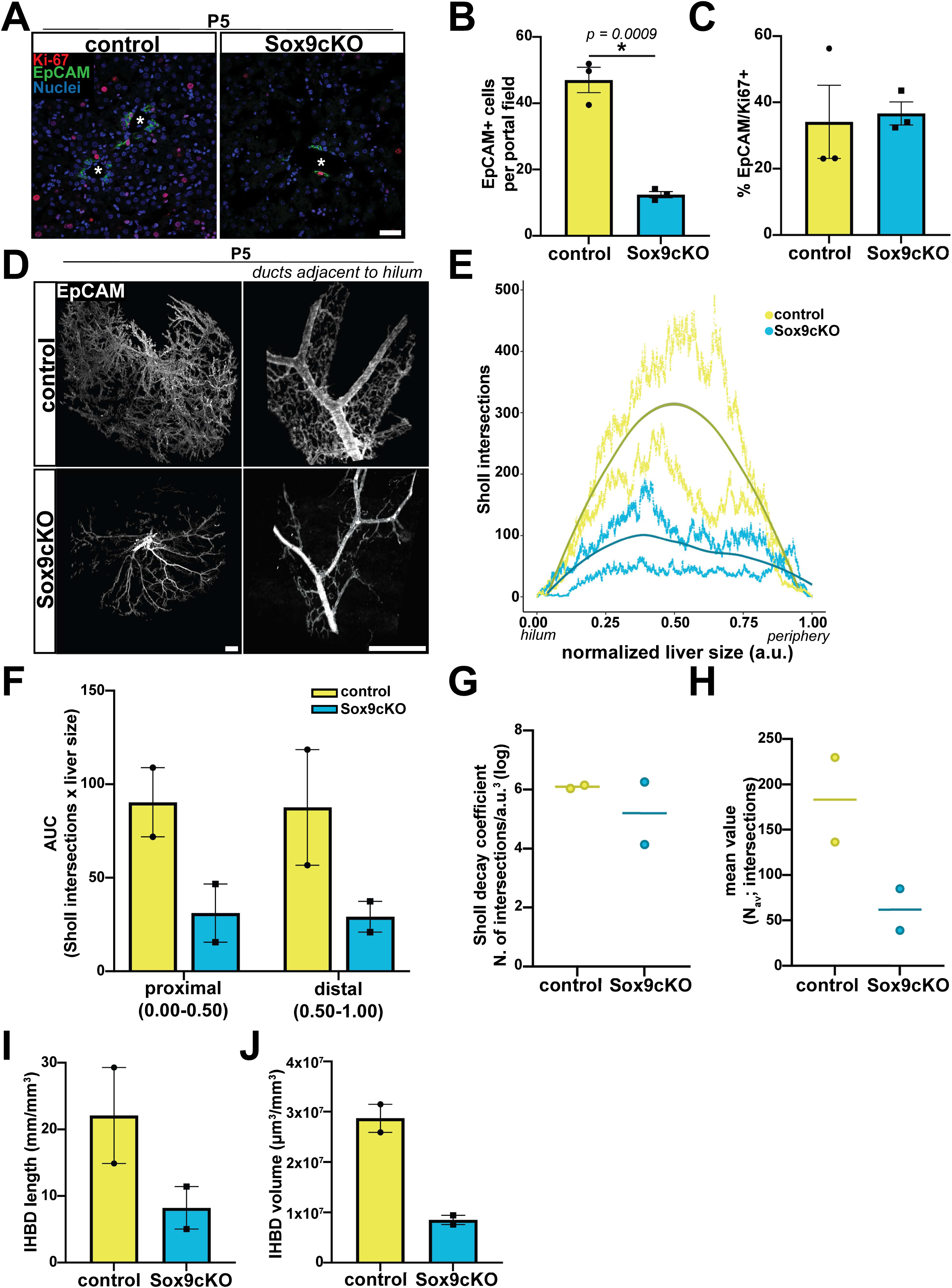
Sox9 regulates branching morphogenesis during IHBD development. **(A)** Co-localization of Ki-67 and EpCAM at P5 shows that Sox9cKO mice exhibit **(B)** significant loss of BECs per portal field, but **(C)** no change in proliferation (scale bar represents 50μm; white asterisks indicate portal vein). **(D)** Whole tissue imaging of P5 livers reveals substantial loss of web-like peripheral IHBDs in Sox9cKO (scale bar represents 0.5mm). **(E)** Sholl analysis demonstrates a loss of branching throughout the liver, which is confirmed by **(F)** AUC showing decreased values in proximal (0.00-0.50a.u.) and distal (0.50-1.00a.u.) regions of Sox9cKO livers. **(G)** Sholl decay coefficient is unchanged between control and Sox9cKO IHBDs, reflecting broad loss of branching versus changes localized to the hilum or periphery. **(H)** The Sholl mean value (N_av_) is also decreased in P5 Sox9cKO mice, as is **(I)** total IHBD length and **(J)** total IHBD volume.

### Inhibition of Activin A partially rescues branching morphogenesis in Sox9cKO livers

We reasoned that Activin A inhibition might be sufficient to rescue Sox9cKO IHBD morphology during development, similar to the phenotypic rescue observed in Sox9cKO mICOs. We treated Sox9cKO pups at P1 and P3 with anti-Activin A (α-ActA) monoclonal antibody, which has been previously shown to neutralize Activin A *in vivo* (Figure 7A) ^47–49^. As a control, a second group of Sox9cKO pups was injected with IgG. Conventional IF for EpCAM demonstrated that Sox9cKO mice treated with α-ActA exhibited increased BEC numbers per portal field when compared to IgG injected Sox9cKO mice (Figure 7B & C). Surprisingly, neutralization of Activin A decreased the number of Ki-67+ BECs, as well as overall liver volume as measured by light sheet imaging (Figure 7B, D, & S6A). These data suggest that Activin A promotes BEC proliferation and liver size, while inhibiting BEC specification during postnatal IHBD development.

**Figure 7.**
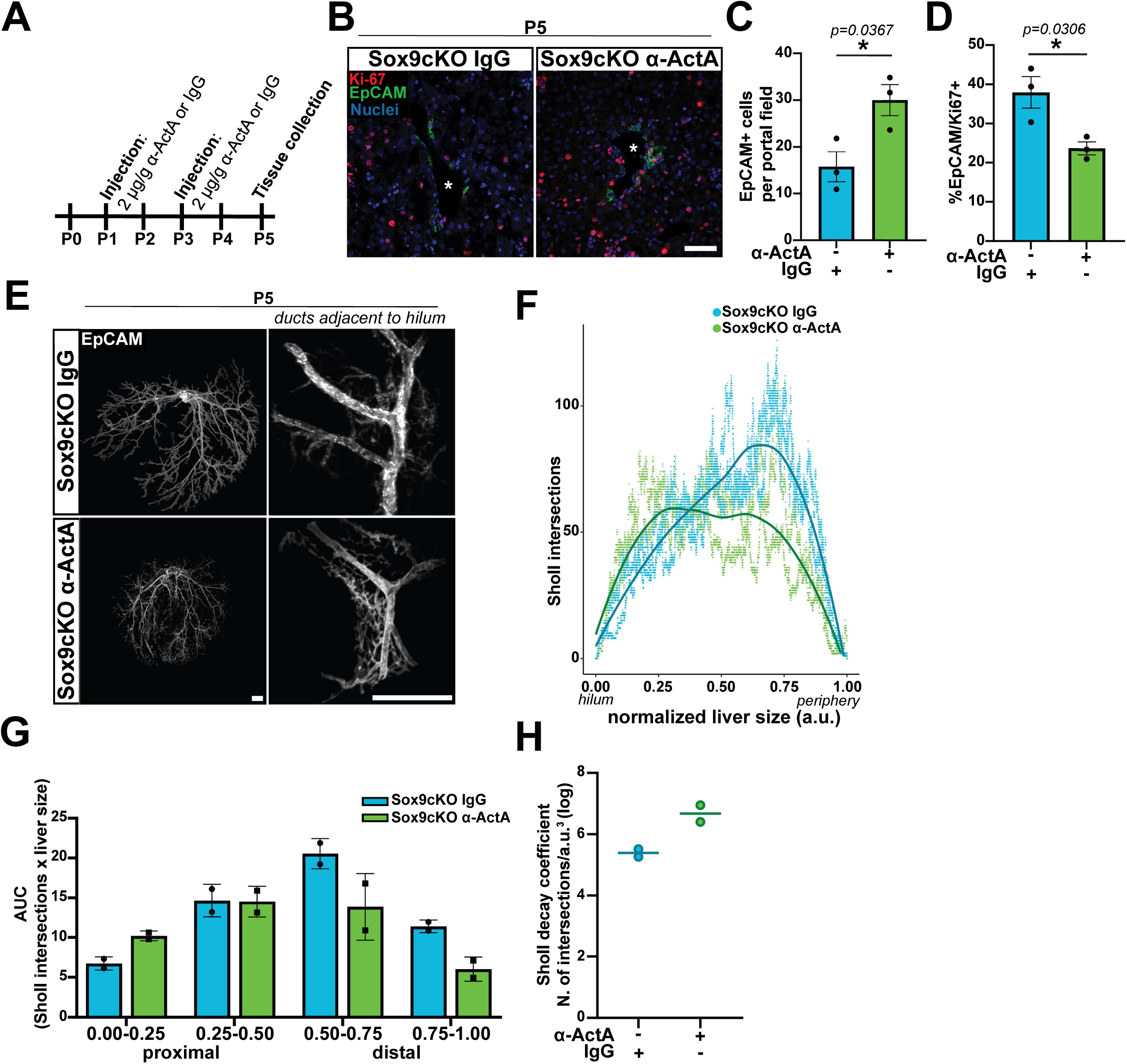
Activin A inhibits IHBD branching morphogenesis during postnatal development. **(A)** Co-localization of Ki-67 and EpCAM in IgG and α-ActA treated livers demonstrates that Activin A neutralization *in vivo* **(B)** increases BEC numbers per portal field while **(C)** inhibiting BEC proliferation (scale bar represents 50μm; white asterisks indicate portal vein). **(D)** Whole tissue imaging of EpCAM+ IHBDs reveals that Activin A neutralization partially rescues Sox9cKO ductal paucity, resulting in increased branching near the hilum. **(E)** Sholl analysis supports increased proximal branching and decreased distal branching in α-ActA treated livers, which is corroborated by **(F)** AUC demonstrating increased hilar branch complexity (0.00-0.25a.u.), no change at 0.25-0.50a.u., and decreased branching distally (0.50-1.00a.u.). **(G)** Sholl decay coefficient is increased following Activin A neutralization reflecting increased localized branching near the hilum.

To determine how Activin A impacts duct vs. ductule morphogenesis, we performed whole-tissue imaging of P5 Sox9cKO livers treated with α-ActA or IgG control antibody. Three dimensional reconstructions of light sheet images revealed a clear web-like IHBD network surrounding main ducts near the hilum of Sox9cKO mice treated with α-ActA antibody, while IgG-treated IHBDs resembled untreated P5 Sox9cKO samples (Figure 7E & Video S4). To quantify this observation, we carried out Sholl analysis, which confirmed increased IHBD branch intersections near the liver hilum in α-ActA treated livers (Figure 7F & S6B). Because the effect of Activin A neutralization was observed in close proximity to the hilum, we carried out AUC analysis on quartiles of normalized Sholl intersections, moving from the hilum to the periphery. AUC was substantially increased in α-ActA treated Sox9cKO livers in the most proximal quartile (0.00-0.25a.u.) and decreased approaching the periphery (0.50-0.75a.u. and 0.75-1.00a.u.) (Figure 7G). This may be related to decreased BEC proliferative expansion, observed in tissue sections. Increased hilar branching in α-ActA treated Sox9cKO livers was further supported by increased Sholl decay coefficient (Figure 7H). No differences was noted between IgG and α-ActA livers for N_av_, Max, r_c_, or total IHBD length and volume, which may be attributed to the shift in branching from the periphery to the hilum following Activin A neutralization (Figure S6C-J). A slight decrease in N_m_ was observed consistent with differences in peripheral branching (Figure 7F & S6D). These data support a critical role for Activin A in postnatal IHBD branching morphogenesis, where it promotes BEC proliferation but limits web-like branching from main ducts.

## Discussion

BEC specification and initial tubulogenesis are well-studied aspects of IHBD branching morphogenesis, but mechanisms underlying IHBD expansion and branch ramification have remained elusive. *Sox9* is one of the earliest markers of BECs in the ductal plate and is commonly associated with BEC identity. Previous studies have shown that *Sox9* regulates the timing of early BEC specification and can worsen ductal paucity in the context of Notch deficiency, but it has been thought to be dispensable for IHBD formation ^10,50^. We show that loss of *Sox9* in hepatoblasts results in a failure to form web-like ductal structures in postnatal IHBD development. Importantly, this manifests as lifelong disorganization of IHBD architecture in adult mice, characterized by reduced numbers of ductules. Results from our P5 and adult imaging of Sox9cKO livers suggest that adult ductules are derived directly from web-like structures surrounding the portal vein during development, and point to distinct genetic requirements for morphogenesis of ducts and ductules.

We also find that *Sox9* represses *Inhba*/Activin A, which is a well-established inhibitor of branching morphogenesis in multiple tissues ^33,34^. BEC specification is known to involve a gradient of TGF-β and Activin A that is highest near the portal vein ^3^. Our analyses show that BECs produce *Inhba*/Activin A, which may contribute to an autocrine or paracrine “niche” that promotes TGF-β signaling and regulates local IHBD architecture. Similar to morphological defects, elevated TGF-β signaling and NCAM1 expression persist in adult Sox9cKO BECs, suggesting that *Sox9* plays a key role in BEC maturation by repressing Activin A signaling in a subpopulation-specific manner. This is further supported by the inability of Sox9cKO BECs to form clearly defined lumens in mICO assays, a phenotype that may be attributable to previously characterized polarity defects in the absence of *Sox9* ^25^. Differentially modulating Activin A signaling is sufficient to: (i) induce morphological defects and *Sox9* and *Ncam1* expression in WT mICOs, and (ii) rescue lumenal phenotypes in Sox9cKO mICOs. These data point to a BEC-autonomous negative feedback loop, where BEC-derived Activin A induces *Sox9*, which in turn represses *Inhba*/Activin A. The ability of Activin A inhibition to rescue hilar branching in postnatal IHBD morphogenesis provides functional context for the *Sox9*-*Inhba* relationship and demonstrates that downregulation of Activin A is sufficient to rescue ductule morphogenesis even in the absence of *Sox9*. This aligns with work in other tissues showing that extended treatment of tissue explants with Activin A can result in a complete loss of branched epithelial networks ^34^. Together these data establish repression of Activin A by *Sox9* as an important event in BEC maturation and IHBD branching morphogenesis.

Technical limitations in whole-tissue imaging of adult and developing livers, including tissue size, density, and opacity, have been long-standing barriers to studying IHBD biology. Models for IHBD development have relied on computational reconstruction of tissue sections or retrograde ink injection of ductal lumens ^12,51^. The latter is advantageous in its ability to reduce technical artifacts associated with tissue sectioning and image reconstruction, but ink injections only label continuous structures, may not perfuse into finer ductules, and are not amenable to quantitative analysis. Our whole-tissue imaging was facilitated by technical advances in tissue clearing and large-volume imaging by light sheet microscopy and allowed direct, sensitive visualization of BECs by IF without physical manipulation of IHBD lumens. This approach also allows us to apply Sholl analyses to quantify global patterns in IHBD branching ^31^. Notably, we observe that pronounced, large diameter ducts are not clearly established until P5, in conflict with earlier reports describing emergence of duct-ductule hierarchy at E17.5 ^12^. These studies relied on ink injection, which may have potentially increased intraluminal forces and, accordingly, the diameter of main branches in the developing ductal web. Surprisingly, we find that IHBDs continue to exhibit highly ramified ductule branches as late as P14, and do not have reduced numbers of ductules surrounding large hilar ducts, as observed in 6-12 week old mice. These studies establish iDISCO+ and light-sheet microscopy as valuable tools for understanding IHBD biology, and underscore the need to expand understanding of postnatal IHDB remodeling.

Our whole-tissue imaging also builds on conventional models of early IHBD morphogenesis and expansion. Newly-specified BECs are thought to form at and expand away from the hilum as discontinuous luminal cysts, interconnecting to form IHBDs ^2,51^. Consistent with this model, we observed rare peripheral EpCAM+ cells at the edges of IHBD webs at E15.5. However, by E17.5 and beyond, all EpCAM+ cells appear integrated into a continuous IHBD network. Because we do not examine earlier timepoints, it is possible that joining of discontinuous cysts is the predominant mechanism for formation of the hilar IHBD web that is apparent by E15.5. Additionally, our studies rely on expression of EpCAM+ as it is highly specific to BECs and amendable to iDISCO+ protocols. However, it remains possible that very early BECs or BEC precursors that do not yet express EpCAM may contribute to IHBD expansion in a discontinuous manner, even at developmental timepoints later than E15.5. The role of BEC proliferation during IHBD morphogenesis has also remained ambiguous. Prior studies have speculated that BECs have minimal contributions to IHBD growth after specification from hepatocytes, implying that IHBD expansion during development is driven primarily by continued differentiation of a progenitor population ^17^. Recent studies using scRNA-seq and chromatin assays have established that the liver is devoid of cells with genetic characteristics of hepatoblasts after E14.5, calling into question how IHBDs continue to expand after this time ^35^. Here we show that pre-existing BECs are highly proliferative throughout embryonic and postnatal development, suggesting that BEC proliferation is critical for IHBD expansion and branching morphogenesis.

SOX9 is known to promote stereotypical branching morphogenesis in other tissues, including lung, thyroid gland, and salivary gland, where it is highly localized to actively advancing or branching tips ^19–22^. One key difference in the liver is that SOX9 is expressed by all intrahepatic BECs ^10^. Our data show that IHBDs undergo complex branching morphogenesis, resembling stochastic branching, to form a *Sox9*-dependent webbed structure of IHBDs that has been described as a biliary plexus ^2^. It is possible that variable local levels of *Sox9* in IHBDs could drive branch formation. Heterogeneous expression of *Sox9* has been reported in the pancreatic ducts, which undergo stochastic branching morphogenesis, with highest *Sox9* expression in small terminal ducts ^52^. Similarly, our lab has previously shown that *Sox9* levels are heterogeneous in IHBDs, with highest *Sox9* levels in ductules ^23^. Therefore, it is possible that high *Sox9* levels drive spatially localized branching programs in parallel with epithelial maturation, in part through suppression of Activin A. Collectively, our studies refine models for IHBD development by identifying distinct genetic requirements for *Sox9*-independent morphogenesis of large ducts and *Sox9*-dependent morphogenesis of small ductules.

## Methods

### RESOURCE AVAILABILITY

#### Lead contact

Further information and requests for resources and reagents should be directed to and will be fulfilled by the lead contact, Adam Gracz

#### Materials availability

This study did not generate any unique reagents and the mouse lines generated in this study are available upon execution of a suitable Materials Transfer Agreement.

#### Data and code availability

All high-throughput sequencing data are available in Gene Expression Omnibus (GEO) under the following accession numbers: bulk RNA-seq: GSE249385; scRNA-seq: GSE 249558.

### Animal Studies

*Sox9^fl/fl^* mice (JAX stock #013106) were crossed to mice carrying the Albumin-Cre (Alb-Cre; JAX stock #003574) allele to establish Sox9cKO (Sox9^fl/fl^: Alb^Cre^) mice. Alb^Cre^ was maintained at heterozygosity. All animals were maintained on a mixed background and adult mice were between 6-12 weeks of age when tissue was collected. Mice of both sexes were used throughout each study. Mice were genotyped by PCR using previously published, allele-specific primers ^53,54^. For liver injury studies, mice were administered with 0.3% (wt/vol) thioacetamide (TAA) *ad libitum* in drinking water for 1 month.

To neutralize Activin A, control and Sox9cKO neonates were administered 2 µg/g anti-Activin A (Biotechne; MAB3381) or mouse IgG_1_ (Biotechne; MAB002) at P1 and P3 by intraperitoneal injection. Injection volumes were 30uL per mouse and antibodies were reconstituted in sterile PBS. Neonates were anesthetized prior to injection by placing neonates in a latex sleeve and immersing the neonate to the neck in wet ice for 2-4min as previously described ^55^. Liver tissue was collected 2 days after the final injection (P5). All animal studies were reviewed and approved by the Institutional Animal Care and Use Committee of Emory University.

### Perfusion and tissue processing

Adult and 2wk animals were sacrificed and tissues fixed by intracardiac perfusion with cold PBS followed by cold 4% paraformaldehyde (PFA; Thermo Fisher; 41678-5000) using a peristaltic pump (Kamoer DIpump550). Embryos and neonatal mice under 10 days of age were not perfused. Instead, livers were dissected, swirled in Dulbecco’s Phosphate-Buffered Saline (DPBS; Thermo Fisher; 14190250) to wash off excess blood, and drop fixed in cold 4% PFA. For all samples, dissected livers were fixed overnight in 4% paraformaldehyde at 4°C. The next day, tissues for whole mount imaging were moved to 0.05% sodium azide (Ricca Chemical; RC7144.8) in PBS at 4°C until further processing. Tissues for conventional histology were moved to 30% sucrose overnight, then embedded in Tissue-Tek Optimal Cutting Temperature media (Sakura Finetek USA, Torrance, CA) and frozen on dry ice. Conventional histology tissues and sections were stored at –80°C until use.

### Intrahepatic bile duct dissociation and FACS

BECs were dissociated as previously described, with minor modifications ^23,45^. Liver lobes were dissected avoiding extrahepatic duct and gallbladder to prevent contamination of intrahepatic biliary preps with extrahepatic biliary tissue. Tissue was rinsed in DPBS then minced to small pieces (<0.5cm^2^) using a razor blade. Sample was transferred to a 50mL conical and suspended in prewarmed 37°C Liver Wash Buffer [LWB; DMEM-H (Fisher Scientific; 11965118), 1% fetal bovine serum (Genesee Scientific; 25-525H), 1% GlutaMAX (Fisher Scientific; 35050061), 1% Pen/Strip (Fisher Scientific; 15140122)]. Tissue was allowed to sediment and then supernatant discarded. Tissue was resuspended in LWB and then discarded after sedimentation 2 more times. The tissue was pelleted in a swinging bucket centrifuge at 4°C and 600g for 5min. Supernatant was discarded and sample was resuspended in room temperature (RT) Liver Digest Buffer [0.15 mg/mL collagenase type XI (Sigma Aldrich; C9407), 0.3 U/mL dispase (Corning; 354235), 200 ug/mL DNase (Sigma Aldrich; DN25)] and incubated at 37°C on a rotator for 90min. Every 30min the sample was retrieved and resuspended by pipetting vigorously with a P1000 micropipette set to 1mL to further promote liver dissociation. After 90min, an equivalent volume of LWB was added to the dissociation and the tissue was pelleted at 4°C and 200g for 5min. The supernatant was carefully decanted to ensure the pellet remained intact and then the sample was washed again with LWB, pelleted, and supernatant decanted. Samples were resuspended in 5mL RBC lysis buffer (BioLegend; 420301) diluted in molecular grade water and incubated on ice for 5min with periodic agitation. RBC Lysis was quenched using 15mL DPBS and the sample was pelleted at 4°C and 600g for 5min and supernatant decanted. For whole duct organoid experiments, dissociation was stopped at this point and duct fragments were used for organoid culture.

To further dissociate liver tissue to single cells for FACS, samples were resuspended with 3mL TrypLE (Fisher Scientific; 12605010) + 10µM Y-27632 (Selleck Chemicals; S1049) and incubated at 37°C for 12min. Every 2min, the sample was pipetted vigorously for 1min with a P1000 micropipette set to 1mL, to mechanically dissociate IHBDs. After 12min, the TrypLE was quenched using 10mL DPBS, and the sample was pelleted at 4°C and 600g for 5min. The supernatant was decanted, and pellet resuspended in Sort Media [Advanced DMEM/F12 (Thermo Fisher; 12634028) with 2% B27 supplement without vitamin A (Thermo Fisher; 12587010), 1% N2 supplement (Thermo Fisher; 17502048), 2mM Glutamax (Thermo Fisher; 35050061), 10mM HEPES (Thermo Fisher; 15630080), 100 U/mL Pen/Strip (Fisher Scientific; 15140122), and 0.1% DNase (Sigma-Aldrich; DN25)]. After thoroughly resuspending, sample was filtered through a 40µm cell strainer (Falcon; 352340).

Single cells were stained with anti-CD31-APC (1:100) (BioLegend; 102510), anti-CD45-APC (1:100) (BioLegend; 103112), and anti-CD326-APC/Cy7 (1:100) (BioLegend; 118218) for 1hr on ice. Cells were rinsed in Sort Media, pelleted at 4°C and 600g for 5min, and supernatant decanted. Cells were resuspended in sort media. 5uL of 7-AAD (BioLegend; 420404) and 5 μL Annexin V-APC (BioLegend; 640941) were added to resuspended cells to distinguish dead and dying cells, respectively. Cells were immediately analyzed and collected on a Sony SH800S fluorescence-activated cell sorter or BD FACSAria II (BD Biosciences). Gating schemes are shown in Figure S2A.

### Mouse intrahepatic biliary organoid culture

8,000 cells per sample were isolated using FACS and collected into 500μL of Sort Media in 1.5 mL polypropylene tubes for biliary organoid culture. Cells were centrifuged at 6,000g for 5min. Pellet was visualized by eye, supernatant was discarded, and cells were resuspended in 1 part Advanced DMEM/F12 and 2 parts Cultrex Reduced Growth Factor Basement Membrane Extract (R&D Systems; 3533-010-02) for a final solution of 66% Cultrex. 96 well plates were prewarmed to 37°C and cell suspensions were plated as 10μL droplets and allowed to polymerize at 37°C for 30min. After polymerization, Biliary Single Cell Media [50% Advanced DMEM/F12, 40% WNT3A-conditioned media, 10% RSPO1-conditioned media, 2% B27 supplement without vitamin A, 1% N2 supplement, 2mM Glutamax, 10mM HEPES, 100 U/mL Pen/Strip, 50ng/mL recombinant murine EGF (Thermo Fisher; PMG8041), 100ng/mL recombinant human Noggin (VWR International; 120-10C), 100ng/mL recombinant human FGF10 (Peprotech; 100-26), 10nM recombinant human gastrin (Sigma; G9145), 5ng/mL recombinant human HGF (PeproTech; 100-39H), 10mM nicotinamide (Sigma Aldrich; N0636-100G), 10µM forskolin (Selleck Chemicals; S2449), and 10µM Y-27632]. Media was changed every other day. For treatments with 50ng/mL Activin A (BioLegend; 592002) or 10μM SB431542 (Selleck Chem; S1067), the reagents were added to the Biliary Single Cell Media for the entire culture period. At the end of 7 days, organoids were analyzed for organoid formation, area, and morphology. Organoid area was measured using bright field images taken using Olympus IX-83 inverted epifluorescent microscope and cellSens Imaging software. Non-cystic morphology was defined as biliary organoids that lack a well-defined lumen. For gene expression analysis, organoids were resuspended in 500μL RNA Lysis Buffer (Invitrogen RNAqueous Micro Kit; AM1931) and lysates were stored at −80°C.

For whole mount organoid experiments, organoids were grown from ductal fragments. Like single cells, ductal fragments were resuspended in 2 parts Cultrex and 1 part Advanced DMEM/F12. Single-cell media was used for the first 2 days in culture following biliary isolation or passaging. After 2 days, the media was changed to Biliary Expansion Media [90% Advanced DMEM/F12, 10% RSPO1-conditioned media, 2% B27 supplement without vitamin A, 1% N2 supplement, 2mM Glutamax, 10mM HEPES, 100 U/mL Pen/Strip, 50ng/mL recombinant murine EGF, 100ng/mL recombinant human FGF10, 10nM recombinant human gastrin, 5ng/mL recombinant human HGF, 10mM nicotinamide, and 10µM forskolin]. Organoids were passaged every 5-7 days.

For statistical comparison, one-way ANOVA was used. A value of *p* < 0.05 was considered significant. Statistical analyses were carried out in Prism 10.1.0. All values are depicted as mean ± SEM.

### Immunofluorescence on tissue sections

10μm tissue sections were cut on a cryostat (Leica; CM1860) and allowed to dry for 4-6hr at RT. Sections were washed 3×5min with PBS, permeabilized for 20min with 0.3% Triton X-100 (Sigma; T8787-250ML) in PBS, and then blocked in 5% normal donkey serum (NDS; Jackson Immuno; 017-000-121) for 45min. If using mouse primary antibody, tissues were then blocked with 1x mouse on mouse (MOM) blocking reagent (Vector Laboratories; MKB-2213) in PBS for 1hr, washed 2×5min with PBS, and rinsed briefly with PBS twice before proceeding with protocol. For all slides, primary antibody was applied overnight at 4°C in PBS. The next day, primary antibody solution was discarded, and slides were rinsed 3×5min with PBS. Secondary antibody raised against host species of the primary antibody was applied in PBS for 1hr at RT. After staining, antibody solution was discarded and bisbenzimide (Sigma; 14530) was applied at 1:1000 in PBS for 10min to label nuclei. Slides were then washed 3×5min with PBS, mounted with ProLong Glass Antifade (Thermo Fisher; P36980), and allowed to cure overnight at RT in the dark. Images were acquired on a laser scanning Nikon A1R HD25 Confocal Microscope in the Emory Integrated Cellular Imaging Core using 40X and 60X oil objectives with working distances of 0.14mm. Acquisition settings were consistent between control and Sox9cKO samples. Statistical analyses were carried out in Prism 10.1.0 (GraphPad Software). For statistical comparison, unpaired *t* test was used. A value of *p* < 0.05 was considered significant. Statistical analyses were carried out in Prism 10.1.0. All values are depicted as mean ± SEM.

#### Antibody usage for immunostaining

Primary antibodies were used at the following concentrations for immunostaining: 2.5μg/mL anti-Activin A (R&D Systems; MAB3381), 0.75μg/mL anti-pSMAD2 (Ser465/467) (Cell Signaling; 3108T), 2.5μg/mL anti-EpCAM/CD326 (BioLegend; 118201), 1μg/mL anti-Ki-67 (Invitrogen; PA5-19462), 7μg/mL anti-Ac-alpha-Tubulin (K40), 0.16μg/mL anti-Cleaved Caspase 3 (D175) (Cell Signaling; 5335S), 0.2μg/mL anti-NCAM1 (Cell Signaling; 99746T), 1μg/mL anti-SOX9 (Millipore; AB5535), 2.5μg/mL anti-EpCAM/CD326-647 (BioLegend; 118212). Secondary antibodies used for immunostaining: 4μg/mL donkey anti-rabbit Alexa Fluor 555 (ThermoFisher; A31572), 4μg/mL donkey anti-rat Alexa Fluor 488 (ThermoFisher; A21208), 4μg/mL donkey anti-mouse Alexa Fluor 555 (ThermoFisher; A31570), and 5μg/mL goat anti-rabbit HRP (ThermoFisher; G21234).

#### Tyramide signal amplification (TSA)

TSA was used for secondary detection of pSMAD2. Slides were washed 3×5min with PBS. During the final wash, peroxidase block was made by combining 1 part H_2_O_2_ (30% H_2_O_2_ stock; Sigma; 216763) and 9 parts methanol (Fisher Scientific; A4134). Slides were submerged in a slide mailer with peroxidase block for 10min at room temperature. After treatment, slides were washed 3×5min with PBS and primary antibody was applied overnight in PBS at 4°C. The next day, primary antibody solution was discarded, and slides were washed 3×5min with PBS. Secondary staining was applied including 5μg/mL goat anti-rabbit HRP (ThermoFisher; G21234) for 1hr. Slides were washed 3×5min in 100mM borate, pH 8.5 (J.T. Baker; 0084-01) supplemented with 0.1% Tween-20 (Thermo Fisher; 85113). During the final wash, the TSA reaction buffer was assembled by adding TSA vivid fluorophore (1:500; Tocris Biosciences; 7526 to 100 mM borate, pH 8.5, 0.1% Tween-20, and 0.003 % H_2_O_2_. Solution was applied to tissue and incubated in the dark for 10min. Slides were then washed 3×5min with 0.1% Tween-20 in PBS to reduce background. Finally, bisbenzimide was applied 1:1000 for 10min. Slides were washed 3×5min with PBS and mounted for imaging as described above.

### Whole mount organoid staining

Whole mount organoids were stained as previously described ^56^. All steps from organoid retrieval through fixation were conducted with tubes and pipette tips pre-coated in 1% (wt/vol) PBS–BSA (Sigma Aldrich; A9647), to prevent organoid loss. After fixation, BSA-coated plastics were not used. Organoids were retrieved from Cultrex by removing culture media and adding 500uL Cultrex Organoid Harvesting Solution (R&D Systems; 3700-100-01) to the wells of a 48 well plate. The plate was incubated on a nutating platform for 45-60min at 4°C. After incubation, organoids were retrieved by resuspending well contents 5-10 times, then transferred to a 15mL tube using a 1mL pipette tip. The tube was filled to 10mL with cold PBS and pelleted at 70g for 3min at 4°C. Organoids were fixed by resuspension in 1mL of 4% PFA for 45min at 4°C and gently resuspended halfway through the incubation. Next, organoids were permeabilized with 10mL of cold 0.1% (vol/vol) PBS-Tween for 10min at 4°C. Organoids were pelleted at 70g for 5min at 4°C and supernatant discarded. Organoids were transferred to a 24-well plate in 200μL of cold (4°C) Organoid Wash Buffer (OWB; 0.1% Triton X-100 and 2mg/mL BSA in PBS) and incubated at 4°C to block the organoids. 200μL of OWB was left in each well for buffer changes to reduce organoid loss. Rabbit anti-Ki-67 primary antibody was added 1:500 in 200μL OWB and added to the pre-existing 200μL in the well for a final antibody concentration of 1:1000 and incubated overnight at 4°C on a nutating platform. The next day, primary antibody was diluted out using 1mL of OWB. Organoids were washed 3×2hr on a nutating platform at 4°C with 1mL OWB buffer changes. Once organoids had settled to the bottom of the plate, 1mL of OWB was removed and donkey anti-rabbit Alexa Fluor 555 was applied at 1:250 in 200uL OWB and added to the pre-existing 200uL OWB for a final concentration of 1:500 and incubated overnight at 4°C on a nutating platform. The next day, bisbenzimide diluted 1:500 in 200μL OWB and added to the 200μL volume of OWB remaining in wells for a final concentration of 1:1000 for 30min. Organoids were washed in 1mL OWB 3×2hr on a nutating platform at 4°C. Next, as much OWB was removed as possible, and organoids were cleared by resuspension in 60% (vol/vol) glycerol and 2.5M fructose in water. For mounting, a barrier pen (Vector Labs; H-4000) and double-sided sticky tape was used to make a thin well to contain the organoids and organoids were mounted with a cover slip. Whole mount organoids were quantified and imaged using Nikon A1R HD25 Confocal Microscope in the Emory Integrated Cellular Imaging Core and Olympus IX-83 inverted fluorescence microscope. Analyses were performed using 1μm step size, z-stacks through the entire organoid, and max-intensity projections in FIJI. Statistical analyses were carried out in Prism 10.1.0. For statistical comparison, unpaired *t* test was used. A value of *p* < 0.05 was considered significant. Statistical analyses were carried out in Prism 10.1.0. All values are depicted as mean ± SEM.

### Whole tissue staining and iDISCO+ clearing

Whole liver IHBD labeling was performed largely in accordance with previously published iDISCO+ protocols ^28^. Transferring of livers between buffers was done using the spoon end of a laboratory spatula to avoid damaging the tissue with tweezers. Perfusion-fixed livers were retrieved from 4°C and washed in PBS for 30min. Samples were then dehydrated by incubating in methanol (Fisher Scientific; A4134) diluted in DI water: 20%, 40%, 60%, 80%, 100%, and 100% for 1hr each. The samples were delipidated by incubating overnight with 66% dichloromethane (DCM; Sigma Aldrich; 270997-1L)/33% methanol at RT on a gyrating rocker. The samples were then washed 2×1hr in 100% methanol and chilled at 4°C before bleaching overnight at 4°C using 5% H_2_O_2_ in methanol. The next day, samples were rehydrated with methanol diluted in DI water: 80%, 60%, 40%, 20%, and PBS for 1hr each at RT. Samples were washed twice with PTx.2 (0.2% TritonX-100 in PBS) for 1 hr.

To permeabilize, samples were incubated overnight at 37°C with 80% PTx.2, 23mg/mL glycine (Millipore Sigma; G7126), and 20% DMSO (Sigma Aldrich; D2650). The next day samples were moved to 0.1% Tween-20, 0.1% TritonX-100, 0.1% deoxycholate (Sigma Aldrich; D2510), and 20% DMSO in PBS overnight at 37°C. Next, the samples were blocked for 2d at 37°C in Blocking Solution [84% PTx.2, 6% NDS (Jackson Immuno; 017-000-121), 10% DMSO]. Samples were stained with unconjugated 2.5µg/mL anti-CD326 (BioLegend; 118201) or anti-CD326-647 conjugated antibody (BioLegend; 118212) diluted in PTwH [0.002% Tween-20 and 4 µg/mL heparin (Sigma; H3393 in PBS]/ 5%DMSO/3% NDS and for 7 days. Samples were washed in PTwH 4-5x until the next day. For samples labeled with unconjugated primary antibody, secondary antibody was applied for 5 days at 37°C in PTwH/ 5%DMSO/3% NDS. Once staining completed, samples were washed in PTwH 4-5x until the next day.

Samples were dehydrated in methanol diluted in DI water: 20%, 40%, 60%, 80%, 100%, and 100% for 1hr each at RT. Samples were delipidated by shaking for 3hr on a gyrating rocker in 66% DCM/33% Methanol at RT. To remove remaining methanol, samples were washed 2×15min with 100% DCM, shaking on a gyrating rocker at RT. To clear the tissue, samples were incubated in dibenzyl ether (DBE; Millipore Sigma; 108014) overnight. Care was taken to ensure the storage tube was filled completely to prevent air from oxidizing the sample. Samples were either imaged in DBE or ethyl cinnamate (ECi; Sigma; 112372). For samples imaged in ECi, they were cleared in DBE for at least 24hr then moved to ECi at least 24hr before imaging. See Table S1 for detailed descriptions of staining, clearing, and imaging conditions for each sample.

Developmental livers were agarose embedded to facilitate sample mounting for imaging. Agarose embedding was carried out prior to dehydration. 1% agarose was prepared by microwaving 1g/L agarose (Genesee Scientific; 20-102GP) in 1X TAE. Agarose was allowed to cool slightly by swirling in wet ice until it was no longer steaming, and samples were embedded by dabbing sample dry with a Kimwipe, placing in an OCT mold, and overlaying with 1% agarose. Care was taken to avoid bubbles in the agarose. Samples cooled for at least 10min at 4°C before proceeding with dehydration.

### Light sheet imaging

Cleared tissue samples were imaged using Miltenyi UltraMicroscope Blaze Lightsheet (Blaze) in the Emory Integrated Cellular Imaging Core (Blaze; ex 640, em 680/30), LaVision BioTec UltraMicroscope II (UMII) in the Biomedical Microscopy Core at the University of Georgia, or UMII in the UNC Microscopy Services Laboratory (ex 640, em 680/30). Whole tissue images using the Miltenyi UltraMicroscope Blaze Lightsheet were acquired using a 1.1x MI PLAN objective (WD ≤ 17mm) or 12× MI PLAN objective (WD ≤ 10.9mm) in ECi. LaVision BioTec UltraMicroscope II whole tissue images were acquired using Olympus MVPLAPO 2X/0.5 objective with a custom LaVision BioTec dipping cap (WD ≤ 10mm) in DBE (detailed information in Table S3). Laser power was adjusted based on intensity of the fluorescent signal. Z-step size for whole tissue images was 5μm. High magnification images of the adult tissue was captured on the UMII using a 6.3× zoom, and the Z-step size was 1μm. High magnification of developmental tissue was captured using the Blaze with a 12× objective, 1x zoom, and 1μm step size.

### Whole tissue image processing and data analysis

3D rendering and video generation for the 3D images was done on high-powered workstations in the Emory Integrated Cellular Imaging Core with 512 GB RAM, NVIDIA graphics card, and 8-64 core processors. We used Imaris (v.10.0 and v.10.1, Bitplane) with the Imaris for Neuroscientists package for 3D visualization and image analyses. Image processing and analyses were conducted by using the surfaces function in Imaris and the Labkit extension or Imaris Machine Learning Training for pixel classification to segment positive staining ^57^. In Labkit, GPU-accelerated segmentation was enabled. Following segmentation, images were masked to set non-segmented regions to 0 pixel intensity and segmented regions to 1000 pixel intensity to create binary images. Binary images were used with filament tracer in Imaris. In filament tracer, the ‘autopath (loops) with Soma and no Spine’ was selected. One starting point was defined at the liver hilum. Multiscale points were used for seed points. Once the filament was defined, 3D Sholl analysis of IHBDs was carried out by concentric 3D spheres being implemented every 1µm around the starting point. The number of intersections at each sphere was measured at 1µm step sizes. For statistical comparison, unpaired *t* test was used. A value of *p* < 0.05 was considered significant. Statistical analyses were carried out in Prism 10.1.0. All values are depicted as mean ± SEM.

#### Sholl analysis

Descriptive statistical analyses of Sholl data acquired in Imaris were performed in R, following a previously described framework (Supplemental File 1) ^31,58^. Sholl distance was normalized to a range of 0 to 1 by dividing each Sholl step size by maximum distance assigned a positive intersectional value in Imaris. To compute the polynomial model of Sholl intersections, we designed an algorithm utilizing the *caret* package to select the best-fit polynomial degree (up to a maximum of 7) using Monte-Carlo Cross-Validation (MCCV) to decrease overfitting ^59^. MCCV was performed for 20 iterations with 80% of the data used as a training set and 20% as a testing set, and the polynomial degree with the lowest average Root Mean Square Error was used to fit a final polynomial model on the entire dataset. The mean value (N_av_), critical value (N_m_), and critical radius (r_c_) metrics were computed from this final polynomial model, as previously described ^31^. From the polynomial model, the mean value indicates the average number of Sholl intersections across the sample, the critical value (maximum of the polynomial model) indicates the maximum number of Sholl intersections, and the critical radius indicates the distance from the hilum where N_m_ occurs ^31^. To calculate Sholl decay, Sholl intersections were log-transformed to fit linear models of the Sholl dataset. R-squared values and Sholl decay coefficients (slope of the linear model multiplied by −1) were calculated for each model. To represent trends between biological replicates in the same experimental group, trendlines for Sholl intersections were calculated with locally estimated scatterplot smoothing (LOESS) regression, shown in Figures 1, 6, and 7 ^60^.

### Bulk RNA-sequencing

#### Sample collection, library construction, and sequencing

25,000 BECs were FACS isolated from 3 control and 3 Sox9cKO adult mice. Cells were sorted directly into 500μL RNA Lysis Buffer. After isolation, lysates were immediately placed on ice and stored at −70°C until RNA purification. RNA purification was carried out using the RNAqueous Micro Kit (Invitrogen; AM1931) following manufacturer instructions. RNA concentrations were assessed using Qubit fluorometer (Q33238; Thermo Fisher Scientific) following the protocol for the Quant-iT Qubit RNA HS Assay Kit (Invitrogen; Q32852). Library construction started from 4.9ng of RNA per sample made using the SMARTer Stranded Total RNA-seq Kit v3-Pico Input Mammalian (Takara; 634486). The quality of libraries was checked with the with the High Sensitivity DNA Reagents Kit (Agilent; 5067-4626) on a 2100 Bioanalyzer and cDNA concentrations were obtained using Qubit BR dsDNA Assay Kit (Fisher Scientific; Q32853). All samples were sequenced by the Illumina Next-seq 550 instrument with paired-end 2×75bp read length and sequenced to a depth of at least 25M reads per sample in the Emory Integrated Genomics Core.

#### Data analysis

Reads were demultiplexed, mapped to the reference genome (GRCm39), and deduplicated based on UMIs using Cogent NGS Analysis Pipeline (CogentAP v2.0). The CogentAP pipeline uses Cutadapt v3.4, STAR v2.7.8a, SAMtools v 0.1.19, bedtools v2.29.2, and Subread v2.0.0 ^61–65^. The gene matrix file was output by the CogentAP pipeline. Differential expression (DE) analysis was done using DESeq2 package v1.36.0 ^66^. The apeglm package v1.18.0 ^67^ was used for effect size estimation and lfcShrink log2 fold-change values. Genes were considered differentially expressed if LFC ≥ 1 and q value < 0.05. The clusterProfiler package v4.7.1 was used to perform Gene Set Enrichment Analysis (GSEA). Gene sets for GSEA analyses have previously been published_35,36,68_.

### Single Cell RNA-sequencing

#### Sample collection, library construction, and sequencing

125,000-135,000 BECs per sample were isolated using FACS, processed, and fixed following the manufacturer’s protocol for whole cells (Parse Biosciences; SB1001). scRNA-seq libraries were prepared using split-pool barcoding as described in the Evercode WT V2 kit (Parse Biosciences; SB2001) according to the manufacturer’s instructions. Libraries were sequenced on one lane of an Illumina NovaSeq S4 flow cell (R1: 74 cycles, i7 index: 6 cycles, R2: 86 cycles, i5 index: 0 cycles) in the Emory Nonhuman Primate Genomics Core.

#### Data processing and quality control

For scRNA-seq data, fastq files corresponding to Sox9 control and cKO samples across all Parse sublibraries were demultiplexed using Parse Biosciences pipeline v0.9.6. Reads were mapped to mouse reference genome GRCm39 and transcripts counted in STAR v2.710a within the Parse pipeline ^62^. Gene expression matrices were loaded into Seurat (v4.1.0) for all data visualization and downstream analysis ^69^. For quality control, we removed cells fulfilling any of these criteria from downstream analyses: less than 500 unique molecular identifiers (UMIs) or less than 200 features. We also removed contaminating endothelial cells based on expression of *Cd31*.

#### Cell clustering and data analysis

Seurat objects of two samples were merged with *merge()* function. Data was normalized with the default Seurat’s LogNormalize method using *NormalizeData()* function. After normalization, variable features were found using *FindVariableFeatures()* and data was scaled with *ScaleData()* function. Clustering was completed by using the RunPCA(), FindNeighbors(),and FindClusters() functions with the first 26 principal components and a clustering resolution of 0.6. Nonlinear dimensionality reduction was performed using the uniform manifold approximation and projection (UMAP) technique with the RunUMAP() function implemented in Seurat package. Cluster specific markers were identified by applying the FindAllMarkers() function with the implemented in Seurat MAST test ^70^. Gene sets enrichment scores were calculated using UCell package v2.1.1 ^71^. The scCustomize package v1.1.1 was used to adjust colors on the FeaturePlots^72^.

To determine the percent of control and Sox9cKO cells in each cluster, cell counts were normalized between control and Sox9cKO, and the percent of control and Sox9cKO cells in each cluster was calculated (Figure S2D). Clusters with < 80% of one sample type are considered mixed clusters. Sox9^EGFP-high^ cells have been previously reported to be enriched in small ductules ^23^. Therefore, to assess differentially expressed genes in ducts and ductules and between control and Sox9cKO tissues, we regrouped samples as control or Sox9cKO ductules if the cluster’s mean enrichment score of Sox9^EGFP-high^ gene signature is > 0.163 (Figure S2F-G).

#### Reanalysis of developing hepatoblast, cholangiocyte, and hepatocyte data

To determine gene signature associated with developing epithelial cell types in the liver, we reanalyzed published RNA-seq ^35^. The count matrix was downloaded from GEO (accession number GSE142089). Differential expression analysis was done using DESeq2. For DE, samples collected at different time points (E12.5-E17.5) were merged by cell type to form three groups: hepatoblasts, cholangiocytes, and hepatocytes. Samples from each cell type “group” were compared against all other samples to identify genes significantly upregulated in a single cell type. Following the differential expression analysis, genes with adjusted *p*-value ≤ 0.01 and log2FoldChange > 0.5 were chosen for calculating enrichment scores.

### RT-qPCR

RNA lysate from sorted single cells or organoids was thawed on ice and purified using the RNAqueous Micro Kit (Invitrogen; AM1931) following manufacturer instructions. RNA concentrations were assessed using Qubit fluorometer (Q33238; Thermo Fisher Scientific) following the protocol for the Quant-iT Qubit RNA HS Assay Kit (Invitrogen; Q32852). cDNA was generated using the iScript cDNA Synthesis Kit (BioRad; 1708891) according to manufacturer guidelines. cDNA was diluted 1:3 in molecular grade water. RT-qPCR was carried out using Taqman probes and SsoAdvanced Universal Probes Supermix (BioRad; 1725284). Relative fold change of expression was carried out using the delta-delta C_T_ method with 18S as the internal reference gene ^73^. Taqman assays in this study are: *18S* (Hs99999901_s1), *Ncam1* (Mm01149710) and *Sox9* (Mm00448840_m1). For statistical comparison, one-way ANOVA was used. A value of *p* < 0.05 was considered significant. Statistical analyses were carried out in Prism 10.1.0. All values are depicted as mean ± SEM.

---

**Video S1: *Light sheet imaging of EpCAM+ BECs demonstrates global ductal paucity in adult Sox9cKO livers.*** Control sample is shown in the left panel; Sox9cKO sample is shown in the right panel.

**Video S2: *High-magnification, 3D rendering of IHBDs highlights loss of peripheral ductules and decreased EpCAM signal.*** Control sample is shown in the left panel; Sox9cKO sample is shown in the right panel.

**Video S3: *Sox9cKO mICOs fail to form lumens and establish cystic morphology.*** Control sample is shown in the left panel; Sox9cKO sample is shown in the right panel. Images acquired every 15min for 15hrs (scale bar represents 50μm).

**Video S4: *Activin A inhibition increases hilar branching in P5 Sox9cKO livers.*** IgG injected Sox9cKO sample is shown in the left panel; α-ActA injected Sox9cKO sample is shown in the right panel.

**Table S1:** *Bulk RNA-seq differential expression analysis.*

**Table S2:** *scRNA-seq differential expression analysis.*

**File S1: *R script for descriptive statistical analysis of Sholl intersections*.**

## Acknowledgments

We thank Dr. Pablo Ariel (UNC) for providing training and technical support in light sheet imaging. We thank Dr. April Reedy (Emory) for training on the Ultramicroscope Blaze and Stoyan Ivanov (Emory) for training in 3D image analysis in Imaris. We thank Dr. Saul Karpen (Emory) for providing Alb-Cre mice and Dr. Scott Magness (UNC) for providing *Sox9^fl/fl^* mice. We thank Drs. Saul Karpen, Paul Dawson, Ken Moberg, Jitendra Thakur, and members of the Gracz Lab for constructive discussions and critical reading of the manuscript.

This study was funded by the NIH/NIDDK under award numbers R01DK132653 (Gracz) and F31DK134199 (Hrncir). Research reported in this publication was supported in part by the Emory Integrated Genomics Core (EIGC) shared resource of Winship Cancer Institute of Emory University and NIH/NCI under award number P30CA138292, by the Emory University Emory Integrated Cellular Imaging Core Facility (RRID:SCR_023534) and NIH under Award Number S10 OD032320-01, the Emory NPRC Genomics Core is supported in part by NIH P51 OD011132, by the North Carolina Biotech Center Institutional Support Grant 2016-IDG-1016, the Biomedical Microscopy Core at the University of Georgia, and the Microscopy Services Laboratory, Department of Pathology and Laboratory Medicine, in part by P30 CA016086 Cancer Center Core Support Grant to the UNC Lineberger Comprehensive Cancer Center.

## REFERENCES

1. Ben-Moshe, S., and Itzkovitz, S. (2019). Spatial heterogeneity in the mammalian liver. Nature Reviews Gastroenterology & Hepatology 16, 395–410. 10.1038/s41575-019-0134-x.

2. Lemaigre, F.P. (2020). Development of the Intrahepatic and Extrahepatic Biliary Tract: A Framework for Understanding Congenital Diseases. Annual Review of Pathology: Mechanisms of Disease 15, 1–22. 10.1146/annurev-pathmechdis-012418-013013.

3. Clotman, F., Jacquemin, P., Plumb-Rudewiez, N., Pierreux, C.E., Van Der Smissen, P., Dietz, H.C., Courtoy, P.J., Rousseau, G.G., and Lemaigre, F.P. (2005). Control of liver cell fate decision by a gradient of TGFβ signaling modulated by Onecut transcription factors. Genes & Development 19, 1849–1854. 10.1101/gad.340305.

4. Geisler, F., Nagl, F., Mazur, P.K., Lee, M., Zimber-Strobl, U., Strobl, L.J., Radtke, F., Schmid, R.M., and Siveke, J.T. (2008). Liver-specific inactivation of Notch2, but not Notch1, compromises intrahepatic bile duct development in mice. Hepatology 48, 607–616.

5. Tchorz, J.S., Kinter, J., Müller, M., Tornillo, L., Heim, M.H., and Bettler, B. (2009). Notch2 signaling promotes biliary epithelial cell fate specification and tubulogenesis during bile duct development in mice. Hepatology 50, 871–879.

6. Zhang, N., Bai, H., David, K.K., Dong, J., Zheng, Y., Cai, J., Giovannini, M., Liu, P., Anders, R.A., and Pan, D. (2010). The Merlin/NF2 tumor suppressor functions through the YAP oncoprotein to regulate tissue homeostasis in mammals. Dev Cell 19, 27–38. 10.1016/j.devcel.2010.06.015.

7. Yang, L., Wang, W.-H., Qiu, W.-L., Guo, Z., Bi, E., and Xu, C.-R. (2017). A single-cell transcriptomic analysis reveals precise pathways and regulatory mechanisms underlying hepatoblast differentiation. Hepatology 66, 1387–1401. 10.1002/hep.29353.

8. Gérard, C., Tys, J., and Lemaigre, F.P. (2017). Gene regulatory networks in differentiation and direct reprogramming of hepatic cells. Seminars in Cell & Developmental Biology 66, 43–50. 10.1016/j.semcdb.2016.12.003.

9. Roskams, T.A., Theise, N.D., Balabaud, C., Bhagat, G., Bhathal, P.S., Bioulac-Sage, P., Brunt, E.M., Crawford, J.M., Crosby, H.A., Desmet, V., et al. (2004). Nomenclature of the finer branches of the biliary tree: Canals, ductules, and ductular reactions in human livers. Hepatology 39, 1739–1745. 10.1002/hep.20130.

10. Antoniou, A., Raynaud, P., Cordi, S., Zong, Y., Tronche, F., Stanger, B.Z., Jacquemin, P., Pierreux, C.E., Clotman, F., and Lemaigre, F.P. (2009). Intrahepatic bile ducts develop according to a new mode of tubulogenesis regulated by the transcription factor SOX9. Gastroenterology 136, 2325–2333.

11. Yang, L., Wang, W.H., Qiu, W.L., Guo, Z., Bi, E., and Xu, C.R. (2017). A single-cell transcriptomic analysis reveals precise pathways and regulatory mechanisms underlying hepatoblast differentiation. Hepatology 66, 1387–1401. 10.1002/hep.29353.

12. Tanimizu, N., Kaneko, K., Itoh, T., Ichinohe, N., Ishii, M., Mizuguchi, T., Hirata, K., Miyajima, A., and Mitaka, T. (2016). Intrahepatic bile ducts are developed through formation of homogeneous continuous luminal network and its dynamic rearrangement in mice. Hepatology 64, 175–188. 10.1002/hep.28521.

13. Roskams, T., and Desmet, V. (2008). Embryology of Extra- and Intrahepatic Bile Ducts, the Ductal plate. The Anatomical Record 291, 628–635. 10.1002/ar.20710.

14. Terada, T., and Nakanuma, Y. (1995). Detection of apoptosis and expression of apoptosis-related proteins during human intrahepatic bile duct development. Am J Pathol 146, 67–74.

15. Wang, S., Sekiguchi, R., Daley, W.P., and Yamada, K.M. (2017). Patterned cell and matrix dynamics in branching morphogenesis. Journal of Cell Biology 216, 559–570. 10.1083/jcb.201610048.

16. Villasenor, A., Chong, D.C., Henkemeyer, M., and Cleaver, O. (2010). Epithelial dynamics of pancreatic branching morphogenesis. Development 137, 4295–4305. 10.1242/dev.052993.

17. Tanimizu, N., Miyajima, A., and Mostov, K.E. (2009). Liver progenitor cells fold up a cell monolayer into a double-layered structure during tubular morphogenesis. Mol Biol Cell 20, 2486–2494. 10.1091/mbc.e08-02-0177.

18. Vestentoft, P.S., Jelnes, P., Hopkinson, B.M., Vainer, B., Møllgård, K., Quistorff, B., and Bisgaard, H.C. (2011). Three-dimensional reconstructions of intrahepatic bile duct tubulogenesis in human liver. BMC Developmental Biology 11, 56. 10.1186/1471-213X-11-56.

19. Rockich, B.E., Hrycaj, S.M., Shih, H.P., Nagy, M.S., Ferguson, M.A.H., Kopp, J.L., Sander, M., Wellik, D.M., and Spence, J.R. (2013). Sox9 plays multiple roles in the lung epithelium during branching morphogenesis. Proceedings of the National Academy of Sciences 110, E4456–E4464. 10.1073/pnas.1311847110.

20. Gonçalves, A.N., Correia-Pinto, J., and Nogueira-Silva, C. (2020). ROBO2 signaling in lung development regulates SOX2/SOX9 balance, branching morphogenesis and is dysregulated in nitrofen-induced congenital diaphragmatic hernia. Respiratory Research 21, 302. 10.1186/s12931-020-01568-w.

21. Chatzeli, L., Gaete, M., and Tucker, A.S. (2017). Fgf10 and Sox9 are essential for the establishment of distal progenitor cells during mouse salivary gland development. Development 144, 2294–2305. 10.1242/dev.146019.

22. Liang, S., Johansson, E., Barila, G., Altschuler, D.L., Fagman, H., and Nilsson, M. (2018). A branching morphogenesis program governs embryonic growth of the thyroid gland. Development 145, dev146829. 10.1242/dev.146829.

23. Tulasi, D.Y., Castaneda, D.M., Wager, K., Hogan, C.B., Alcedo, K.P., Raab, J.R., and Gracz, A.D. (2021). Sox9(EGFP) Defines Biliary Epithelial Heterogeneity Downstream of Yap Activity. Cell Mol Gastroenterol Hepatol 11, 1437–1462. 10.1016/j.jcmgh.2021.01.009.

24. Poncy, A., Antoniou, A., Cordi, S., Pierreux, C.E., Jacquemin, P., and Lemaigre, F.P. (2015). Transcription factors SOX4 and SOX9 cooperatively control development of bile ducts. Developmental Biology 404, 136–148. 10.1016/j.ydbio.2015.05.012.

25. Xu, W.P., Cui, Y.L., Chen, L.L., Ding, K., Ding, C.H., Chen, F., Zhang, X., and Xie, W.F. (2021). Deletion of *<scp>Sox</scp>9*in the liver leads to hepatic cystogenesis in mice by transcriptionally downregulating *<scp>Sec</scp>63*. The Journal of Pathology. 10.1002/path.5636.

26. Postic, C., Shiota, M., Niswender, K.D., Jetton, T.L., Chen, Y., Moates, J.M., Shelton, K.D., Lindner, J., Cherrington, A.D., and Magnuson, M.A. (1999). Dual Roles for Glucokinase in Glucose Homeostasis as Determined by Liver and Pancreatic β Cell-specific Gene Knock-outs Using Cre Recombinase. Journal of Biological Chemistry 274, 305–315. 10.1074/jbc.274.1.305.

27. Kamimoto, K., Kaneko, K., Kok, C.Y.-Y., Okada, H., Miyajima, A., and Itoh, T. (2016). Heterogeneity and stochastic growth regulation of biliary epithelial cells dictate dynamic epithelial tissue remodeling. eLife 5, e15034. 10.7554/eLife.15034.

28. Renier, N., Wu, Z., David, Yang, J., Ariel, P., and Tessier-Lavigne, M. (2014). iDISCO: A Simple, Rapid Method to Immunolabel Large Tissue Samples for Volume Imaging. Cell 159, 896–910. 10.1016/j.cell.2014.10.010.

29. Sholl, D.A. (1953). Dendritic organization in the neurons of the visual and motor cortices of the cat. J Anat 87, 387–406.

30. Ristanović, D., Milošević, N.T., and Štulić, V. (2006). Application of modified Sholl analysis to neuronal dendritic arborization of the cat spinal cord. Journal of neuroscience methods 158, 212–218.

31. Ferreira, T.A., Blackman, A.V., Oyrer, J., Jayabal, S., Chung, A.J., Watt, A.J., Sjöström, P.J., and van Meyel, D.J. (2014). Neuronal morphometry directly from bitmap images. Nature Methods 11, 982–984. 10.1038/nmeth.3125.

32. Stanko, J.P., Easterling, M.R., and Fenton, S.E. (2015). Application of Sholl analysis to quantify changes in growth and development in rat mammary gland whole mounts. Reprod Toxicol 54, 129–135. 10.1016/j.reprotox.2014.11.004.

33. Cancilla, B., Jarred, R.A., Wang, H., Mellor, S.L., Cunha, G.R., and Risbridger, G.P. (2001). Regulation of Prostate Branching Morphogenesis by Activin A and Follistatin. Developmental Biology 237, 145–158. 10.1006/dbio.2001.0364.

34. Ritvos, O., Tuuri, T., Erämaa, M., Sainio, K., Hildén, K., Saxén, L., and Gilbert, S.F. (1995). Activin disrupts epithelial branching morphogenesis in developing glandular organs of the mouse. Mechanisms of Development 50, 229–245. 10.1016/0925-4773(94)00342-K.

35. Yang, L., Wang, X., Yu, X.-X., Yang, L., Zhou, B.-C., Yang, J., and Xu, C.-R. (2023). The default and directed pathways of hepatoblast differentiation involve distinct epigenomic mechanisms. Developmental Cell 58, 1688–1700.e1686. 10.1016/j.devcel.2023.07.002.

36. Schaub, J.R., Huppert, K.A., Kurial, S.N.T., Hsu, B.Y., Cast, A.E., Donnelly, B., Karns, R.A., Chen, F., Rezvani, M., Luu, H.Y., et al. (2018). De novo formation of the biliary system by TGFβ-mediated hepatocyte transdifferentiation. Nature 557, 247–251. 10.1038/s41586-018-0075-5.

37. Fabris, L., Strazzabosco, M., Crosby, H.A., Ballardini, G., Hubscher, S.G., Kelly, D.A., Neuberger, J.M., Strain, A.J., and Joplin, R. (2000). Characterization and isolation of ductular cells coexpressing neural cell adhesion molecule and Bcl-2 from primary cholangiopathies and ductal plate malformations. Am J Pathol 156, 1599–1612. 10.1016/s0002-9440(10)65032-8.

38. Francis, H., Glaser, S., Demorrow, S., Gaudio, E., Ueno, Y., Venter, J., Dostal, D., Onori, P., Franchitto, A., Marzioni, M., et al. (2008). Small mouse cholangiocytes proliferate in response to H1 histamine receptor stimulation by activation of the IP3/CaMK I/CREB pathway. Am J Physiol Cell Physiol 295, C499–513. 10.1152/ajpcell.00369.2007.

39. Renzi, A., Glaser, S., Demorrow, S., Mancinelli, R., Meng, F., Franchitto, A., Venter, J., White, M., Francis, H., Han, Y., et al. (2011). Melatonin inhibits cholangiocyte hyperplasia in cholestatic rats by interaction with MT1 but not MT2 melatonin receptors. Am J Physiol Gastrointest Liver Physiol 301, G634–643. 10.1152/ajpgi.00206.2011.

40. Alpini, G., Franchitto, A., Demorrow, S., Onori, P., Gaudio, E., Wise, C., Francis, H., Venter, J., Kopriva, S., Mancinelli, R., et al. (2011). Activation of alpha(1)-adrenergic receptors stimulate the growth of small mouse cholangiocytes via calcium-dependent activation of nuclear factor of activated T cells 2 and specificity protein 1. Hepatology 53, 628–639. 10.1002/hep.24041.

41. Luo, Z.L., Cheng, L., Wang, T., Tang, L.J., Tian, F.Z., Xiang, K., and Cui, L. (2019). Bile Acid Transporters Are Expressed and Heterogeneously Distributed in Rat Bile Ducts. Gut Liver 13, 569–575. 10.5009/gnl18265.

42. Alpini, G., Ulrich, C., Roberts, S., Phillips, J.O., Ueno, Y., Podila, P.V., Colegio, O., LeSage, G.D., Miller, L.J., and LaRusso, N.F. (1997). Molecular and functional heterogeneity of cholangiocytes from rat liver after bile duct ligation. Am J Physiol 272, G289–297. 10.1152/ajpgi.1997.272.2.G289.

43. Alpini, G., Glaser, S., Robertson, W., Rodgers, R.E., Phinizy, J.L., Lasater, J., and LeSage, G.D. (1997). Large but not small intrahepatic bile ducts are involved in secretin-regulated ductal bile secretion. Am J Physiol 272, G1064–1074. 10.1152/ajpgi.1997.272.5.G1064.

44. Marsee, A., Roos, F.J.M., Verstegen, M.M.A., Consortium, H.P.B.O., Gehart, H., de Koning, E., Lemaigre, F., Forbes, S.J., Peng, W.C., Huch, M., et al. (2021). Building consensus on definition and nomenclature of hepatic, pancreatic, and biliary organoids. Cell Stem Cell 28, 816–832. 10.1016/j.stem.2021.04.005.

45. Broutier, L., Andersson-Rolf, A., Hindley, C.J., Boj, S.F., Clevers, H., Koo, B.-K., and Huch, M. (2016). Culture and establishment of self-renewing human and mouse adult liver and pancreas 3D organoids and their genetic manipulation. Nature Protocols 11, 1724–1743. 10.1038/nprot.2016.097.

46. Laping, N.J., Grygielko, E., Mathur, A., Butter, S., Bomberger, J., Tweed, C., Martin, W., Fornwald, J., Lehr, R., Harling, J., et al. (2002). Inhibition of transforming growth factor (TGF)-beta1-induced extracellular matrix with a novel inhibitor of the TGF-beta type I receptor kinase activity: SB-431542. Mol Pharmacol 62, 58–64. 10.1124/mol.62.1.58.

47. Mancinelli, G., Torres, C., Krett, N., Bauer, J., Castellanos, K., McKinney, R., Dawson, D., Guzman, G., Hwang, R., Grimaldo, S., et al. (2021). Role of stromal activin A in human pancreatic cancer and metastasis in mice. Sci Rep 11, 7986. 10.1038/s41598-021-87213-y.

48. Zhang, Z., Wang, J., Chen, Y., Suo, L., Chen, H., Zhu, L., Wan, G., and Han, X. (2019). Activin a promotes myofibroblast differentiation of endometrial mesenchymal stem cells via STAT3-dependent Smad/CTGF pathway. Cell Communication and Signaling 17, 45. 10.1186/s12964-019-0361-3.

49. Spottiswoode, N., Armitage Andrew, E., Williams Andrew, R., Fyfe Alex, J., Biswas, S., Hodgson Susanne, H., Llewellyn, D., Choudhary, P., Draper Simon, J., Duffy Patrick, E., and Drakesmith, H. (2017). Role of Activins in Hepcidin Regulation during Malaria. Infection and Immunity 85, 10.1128/iai.00191-00117. 10.1128/iai.00191-17.

50. Thakurdas, S.M., Lopez, M.F., Kakuda, S., Fernandez-Valdivia, R., Zarrin-Khameh, N., Haltiwanger, R.S., and Jafar-Nejad, H. (2016). Jagged1 heterozygosity in mice results in a congenital cholangiopathy which is reversed by concomitant deletion of one copy of Poglut1 (Rumi). Hepatology 63, 550–565. 10.1002/hep.28024.

51. Takashima, Y., Terada, M., Kawabata, M., and Suzuki, A. (2015). Dynamic three-dimensional morphogenesis of intrahepatic bile ducts in mouse liver development. Hepatology 61, 1003–1011. 10.1002/hep.27436.

52. Rezanejad, H., Ouziel-Yahalom, L., Keyzer, C.A., Sullivan, B.A., Hollister-Lock, J., Li, W.C., Guo, L., Deng, S., Lei, J., Markmann, J., and Bonner-Weir, S. (2018). Heterogeneity of SOX9 and HNF1β in Pancreatic Ducts Is Dynamic. Stem Cell Reports 10, 725–738. 10.1016/j.stemcr.2018.01.028.

53. Postic, C., Shiota, M., Niswender, K.D., Jetton, T.L., Chen, Y., Moates, J.M., Shelton, K.D., Lindner, J., Cherrington, A.D., and Magnuson, M.A. (1999). Dual roles for glucokinase in glucose homeostasis as determined by liver and pancreatic beta cell-specific gene knock-outs using Cre recombinase. J Biol Chem 274, 305–315. 10.1074/jbc.274.1.305.

54. Akiyama, H., Chaboissier, M.C., Martin, J.F., Schedl, A., and de Crombrugghe, B. (2002). The transcription factor Sox9 has essential roles in successive steps of the chondrocyte differentiation pathway and is required for expression of Sox5 and Sox6. Genes Dev 16, 2813–2828. 10.1101/gad.1017802.

55. Navarro, K.L., Huss, M., Smith, J.C., Sharp, P., Marx, J.O., and Pacharinsak, C. (2021). Mouse Anesthesia: The Art and Science. ILAR Journal 62, 238–273. 10.1093/ilar/ilab016.

56. Dekkers, J.F., Alieva, M., Wellens, L.M., Ariese, H.C.R., Jamieson, P.R., Vonk, A.M., Amatngalim, G.D., Hu, H., Oost, K.C., Snippert, H.J.G., et al. (2019). High-resolution 3D imaging of fixed and cleared organoids. Nature Protocols 14, 1756–1771. 10.1038/s41596-019-0160-8.

57. Arzt, M., Deschamps, J., Schmied, C., Pietzsch, T., Schmidt, D., Tomancak, P., Haase, R., and Jug, F. (2022). LABKIT: Labeling and Segmentation Toolkit for Big Image Data. Frontiers in Computer Science 4. 10.3389/fcomp.2022.777728.

58. Team, R.C. (2023). R: A Language and Environment for Statistical Computing_. R Foundation for Statistical Computing.

59. Kuhn, M. (2008). Building Predictive Models in R Using the caret Package. Journal of Statistical Software 28, 1–26. 10.18637/jss.v028.i05.

60. Cleveland, W.S. (1979). Robust Locally Weighted Regression and Smoothing Scatterplots. Journal of the American Statistical Association 74, 829–836. 10.1080/01621459.1979.10481038.

61. Martin, M. (2011). Cutadapt removes adapter sequences from high-throughput sequencing reads. EMBnet.journal 17, 10.10.14806/ej.17.1.200.

62. Dobin, A., Davis, C.A., Schlesinger, F., Drenkow, J., Zaleski, C., Jha, S., Batut, P., Chaisson, M., and Gingeras, T.R. (2013). STAR: ultrafast universal RNA-seq aligner. Bioinformatics 29, 15–21. 10.1093/bioinformatics/bts635.

63. Danecek, P., Bonfield, J.K., Liddle, J., Marshall, J., Ohan, V., Pollard, M.O., Whitwham, A., Keane, T., McCarthy, S.A., Davies, R.M., and Li, H. (2021). Twelve years of SAMtools and BCFtools. GigaScience 10, giab008. 10.1093/gigascience/giab008.

64. Quinlan, A.R., and Hall, I.M. (2010). BEDTools: a flexible suite of utilities for comparing genomic features. Bioinformatics 26, 841–842. 10.1093/bioinformatics/btq033.

65. Liao, Y., Smyth, G.K., and Shi, W. (2014). featureCounts: an efficient general purpose program for assigning sequence reads to genomic features. Bioinformatics 30, 923–930. 10.1093/bioinformatics/btt656.

66. Love, M.I., Huber, W., and Anders, S. (2014). Moderated estimation of fold change and dispersion for RNA-seq data with DESeq2. Genome Biol 15, 550. 10.1186/s13059-014-0550-8.

67. Zhu, A., Ibrahim, J.G., and Love, M.I. (2019). Heavy-tailed prior distributions for sequence count data: removing the noise and preserving large differences. Bioinformatics 35, 2084–2092. 10.1093/bioinformatics/bty895.

68. Coulouarn, C., Factor, V.M., and Thorgeirsson, S.S. (2008). Transforming growth factor-β gene expression signature in mouse hepatocytes predicts clinical outcome in human cancer. Hepatology 47, 2059–2067. 10.1002/hep.22283.

69. Hao, Y., Hao, S., Andersen-Nissen, E., Mauck, W.M., 3rd, Zheng, S., Butler, A., Lee, M.J., Wilk, A.J., Darby, C., Zager, M., et al. (2021). Integrated analysis of multimodal single-cell data. Cell 184, 3573–3587 e3529. 10.1016/j.cell.2021.04.048.

70. Finak, G., McDavid, A., Yajima, M., Deng, J., Gersuk, V., Shalek, A.K., Slichter, C.K., Miller, H.W., McElrath, M.J., Prlic, M., et al. (2015). MAST: a flexible statistical framework for assessing transcriptional changes and characterizing heterogeneity in single-cell RNA sequencing data. Genome Biol 16, 278. 10.1186/s13059-015-0844-5.

71. Andreatta, M., and Carmona, S.J. (2021). UCell: Robust and scalable single-cell gene signature scoring. Comput Struct Biotechnol J 19, 3796–3798. 10.1016/j.csbj.2021.06.043.

72. Marsh, S.E. (2021). scCustomize: Custom Visualizations & Functions for Streamlined Analyses of Single Cell Sequencing.

73. Pfaffl, M.W. (2001). A new mathematical model for relative quantification in real-time RT–PCR. Nucleic Acids Research 29, e45–e45. 10.1093/nar/29.9.e45.

