## Supplemental figure legends for "*Sox9* links biliary maturation to branching morphogenesis"

**Supplemental Figures**

**Figure S1.** ***Impact of Sox9 on BECs and IHBD branch complexity.*** **(A)** Co-localization of Ki-67 and EpCAM during adult homeostasis reveals proliferation is unchanged between control and Sox9cKO (scale bar represents 50µm; white asterisks indicate portal vein). **(B)** Sox9cKO BECs have a loss of Ac-tubulin staining indicating loss of primary cilia (scale bar represents 50µm; white asterisks indicate portal vein). **(C)** Co-localization of Ki-67 and EpCAM indicates BEC proliferation induced by 4 weeks of TAA is unchanged between control and Sox9cKO tissues (scale bar represents 50µm; white asterisks indicate portal vein). **(D)** Sholl analyses (linear and semi-log plots) of individual adult control and Sox9cKO replicates demonstrate ductal paucity which is worse peripherally (arbitrary units, a.u., represent distance normalized to lobe size). **(E)** Sox9cKO maximum branching [max raw value and critical value ($N_m$)] is significantly decreased relative to control. **(F)** Critical radius ($r_c$) is unchanged between groups.

**Figure S2.** ***Transcriptomic analysis of BECs in control and Sox9cKO mice.*** **(A)** EpCAM+ cells were FACS isolated excluding CD31, CD45, 7-AAD, and annexin V positive cells. **(B)** Principal component analysis demonstrates distinct grouping of control and Sox9cKO BEC bulk RNA-seq. **(C)** Heatmap of the top 50 upregulated genes in Sox9cKO BECs includes several involved in TGF-β signaling. **(D)** Targeted analysis of TGF-β ligand expression in bulk RNA-seq reveals that the only significantly upregulated ligand is *Inhba* (TGF-β ligands that are not detected are not shown: *Bmp2*, *Bmp15*, *Nodal*, *Gdf3*, and *Gdf8*). **(E)** Previously published RNA-seq data reveals that *Inhba* is expressed at low levels in hepatoblasts and significantly increases in hepatocytes and BECs over the course of embryonic development (E12.5 – E17.5).

**Figure S3.** ***Sox9 regulates gene expression in ducts and ductules.*** **(A)** Control and Sox9cKO BECs can be classified into 11 transcriptomically distinct clusters. **(B)** We classified subpopulations with < 80% of control or Sox9cKO cells as "mixed" clusters: clusters 0, 1, 2, and 7 are control clusters; clusters 4, 5, 6, and 9 are Sox9cKO clusters; and clusters 3, 8, and 10 are mixed clusters. **(C)** Previously reported small and large duct markers are not enriched in specific BEC subpopulations. **(D)** $Sox9^{EGFP\text{-}high}$ gene signature, known to be enriched in small ductules, is enriched in clusters 1, 2, 6, and 8 by UCell GSEA (enrichment threshold > 0.15). **(E)** UMAP demonstrates re-classification of scRNA-seq data according to duct vs. ductule identity as based on $Sox9^{EGFP\text{-}high}$ gene set enrichment. **(F)** Total number of DEGs between duct BECs (n = 1,860) and ductule BECs (n = 1,830) are unchanged.

**Figure S4.** ***Loss of Sox9 does not impact biliary organoid proliferation.*** **(A)** Ki-67+ nuclei are unchanged between control and Sox9cKO mICOs (scale bar represents 50µm).

**Figure S5.** ***Sox9 promotes IHBD branching morphogenesis in postnatal liver development.*** **(A)** Sholl analyses (linear and semi-log plots) of individual P5 control and Sox9cKO replicates demonstrate broad Sox9cKO ductal paucity (arbitrary units, a.u., represent distance normalized to liver size). **(B)** The maximum branching complexity (Max and $N_m$) in P5 Sox9cKO IHBDs is decreased compared to P5 control. **(C)** The critical radius ($r_c$) is unchanged between groups.

**Figure S6.** ***Activin A neutralization increases branching near the hilum in Sox9cKO IHBDs.*** **(A)** Total liver volume is increased in P5 Sox9cKO IgG and decreased in P5 Sox9cKO α-ActA samples relative to uninjected mice. **(B)** Sholl analyses (linear and semi-log plots) of individual P5 Sox9cKO IgG and α-ActA replicates demonstrate increased branching near the liver hilum in P5 Sox9cKO α-ActA samples. Sox9cKO IgG samples demonstrate greater peripheral branching (arbitrary units, a.u., represent distance normalized to liver size). **(C)** The mean value ($N_{av}$), **(D)** max raw value (Max), **(E)** critical radius ($r_c$), **(I)** IHBD length, and **(J)** IHBD volume are unchanged between groups. **(D)** The critical value ($N_m$) trends down in P5 Sox9cKO treated with α-ActA.

**Table S3:** ***Light sheet imaging parameters.***
