## Supplementary figures and images for "*Sox9* links biliary maturation to branching morphogenesis"

### Figure S1

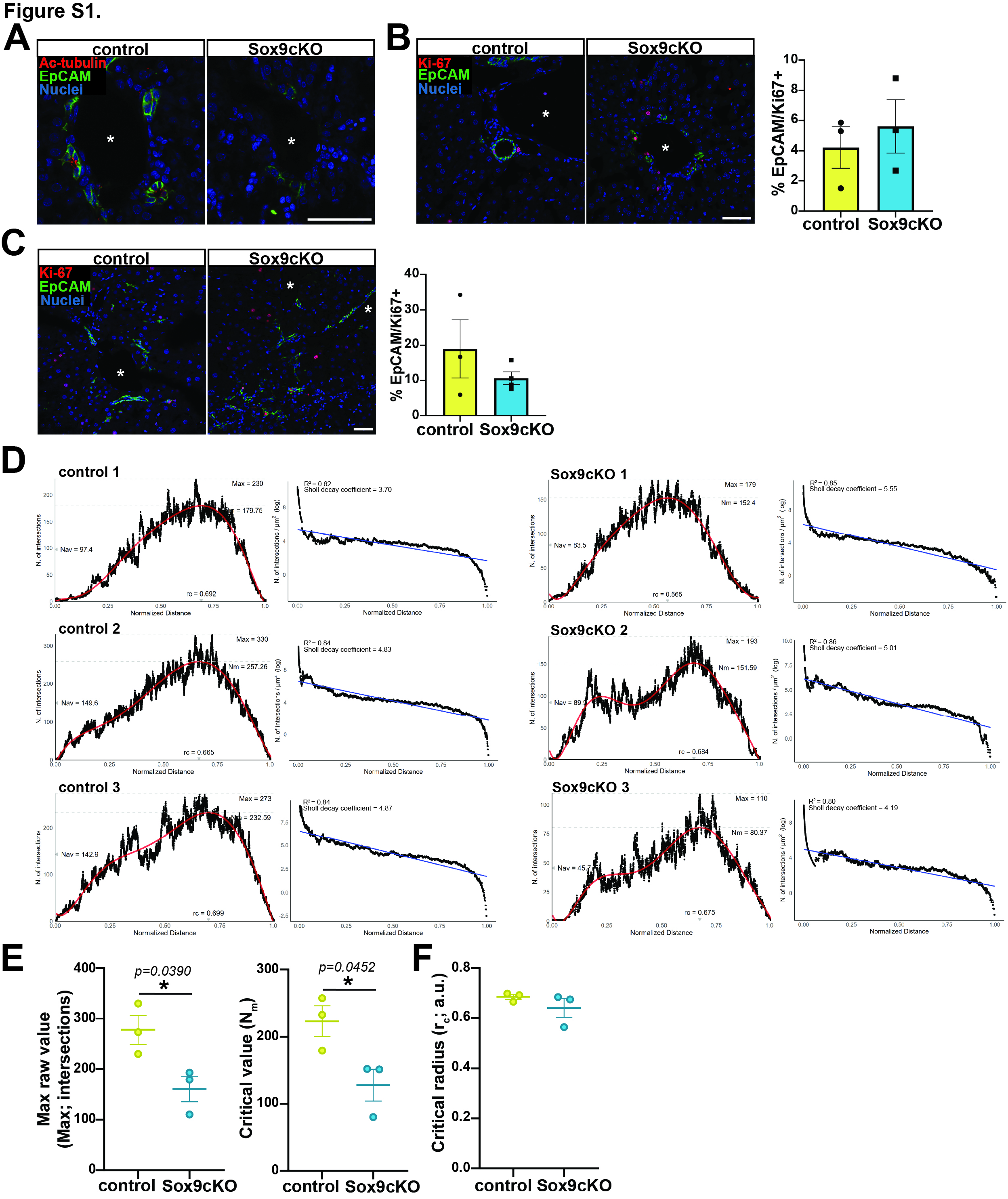

### Figure S2

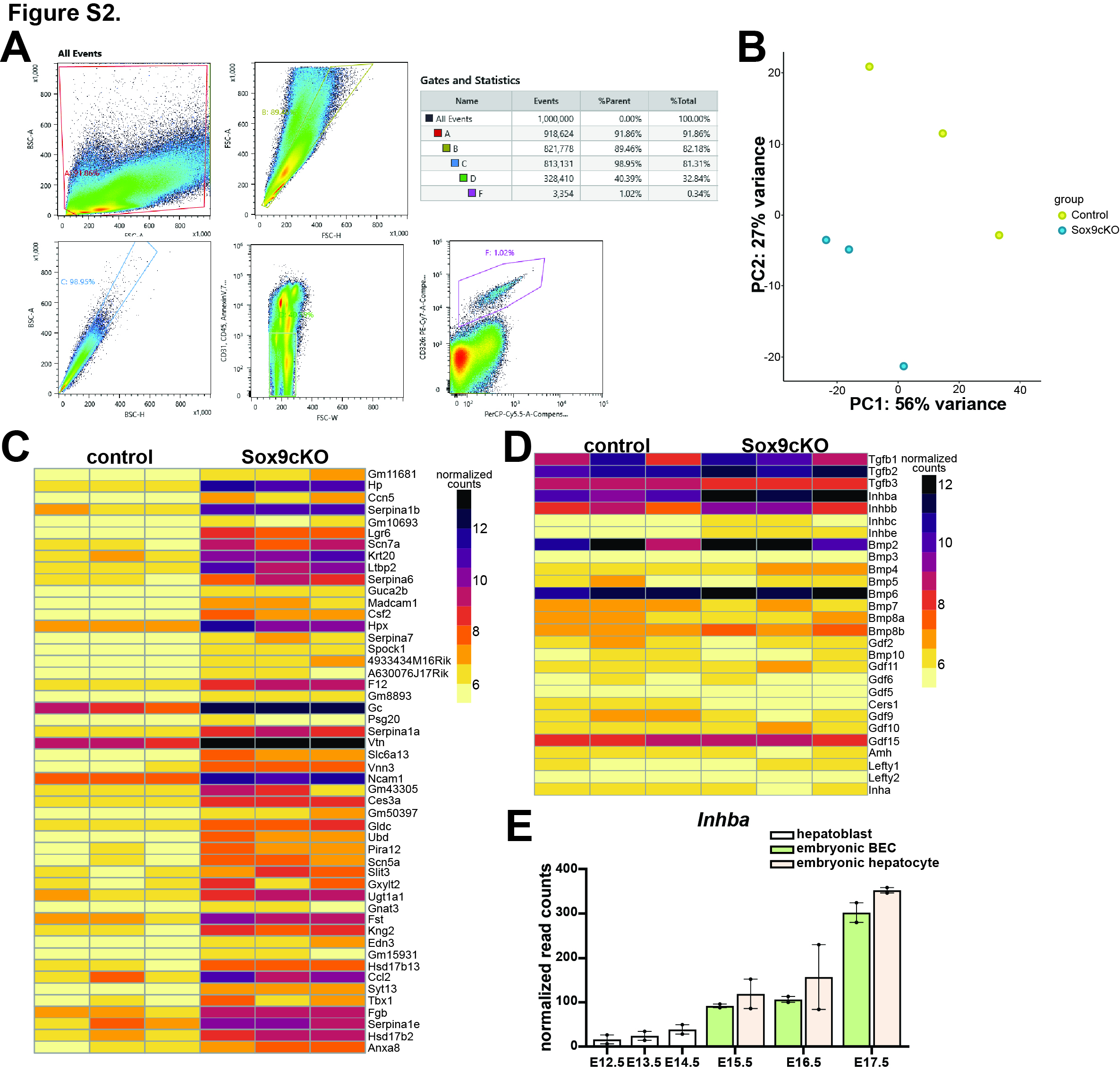

### Figure S3

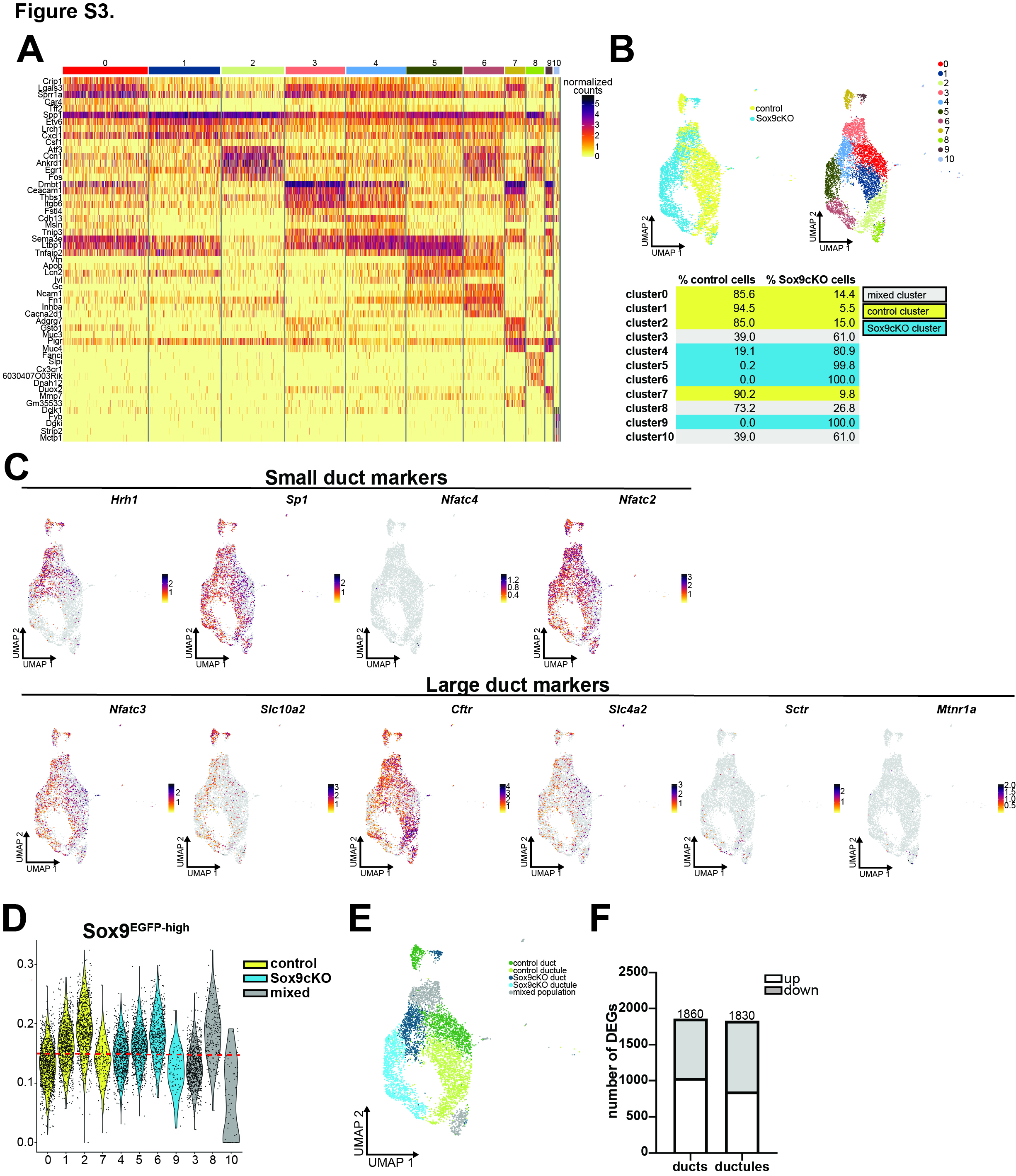

### Figure S4

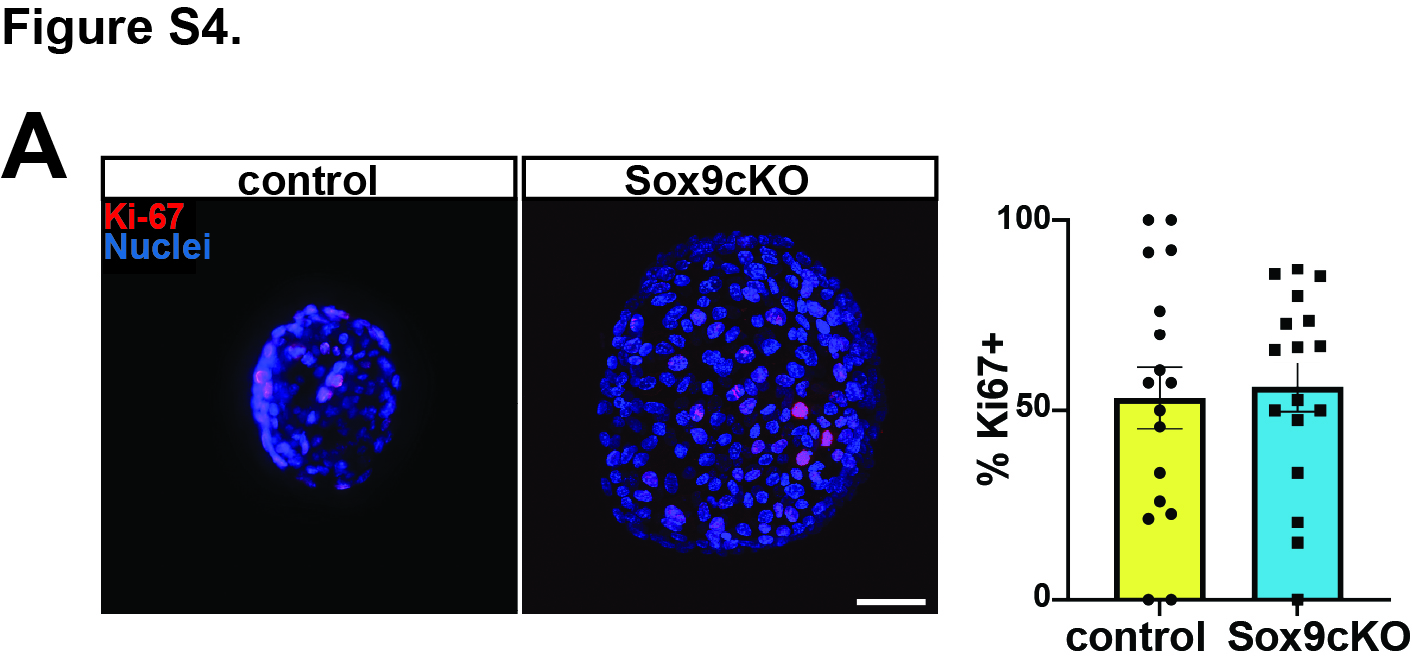

### Figure S5

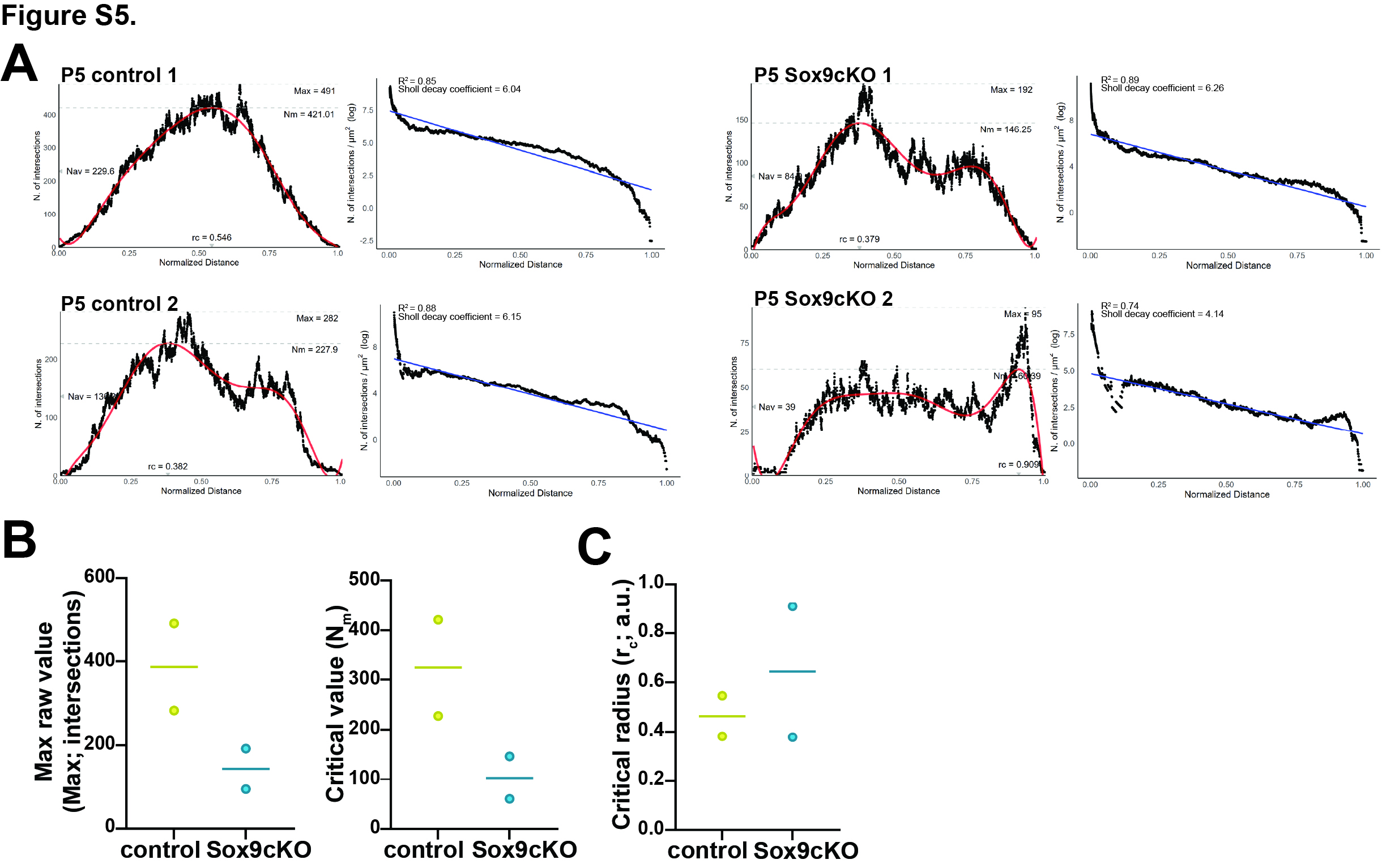

### Figure S6

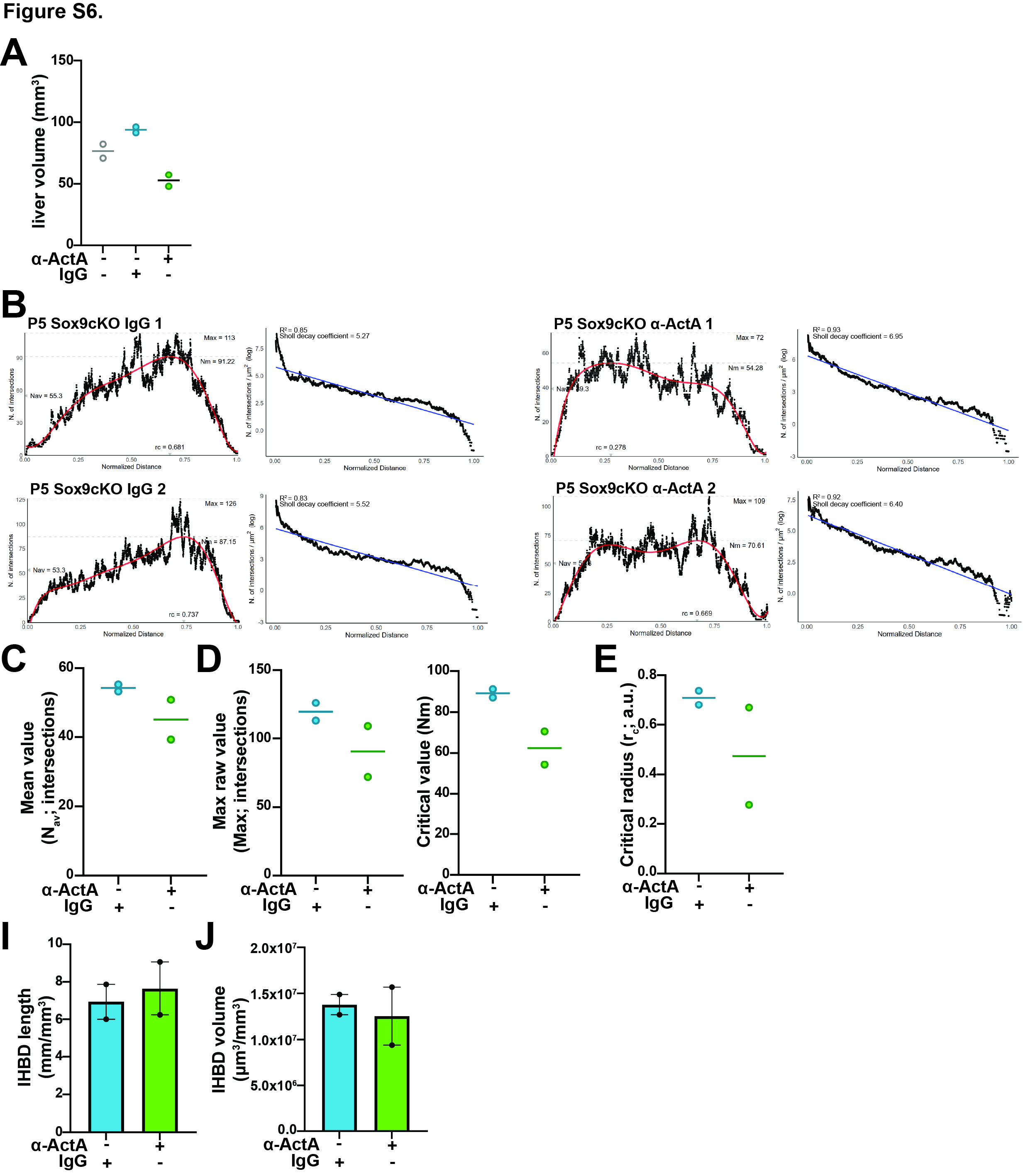

### Table S3

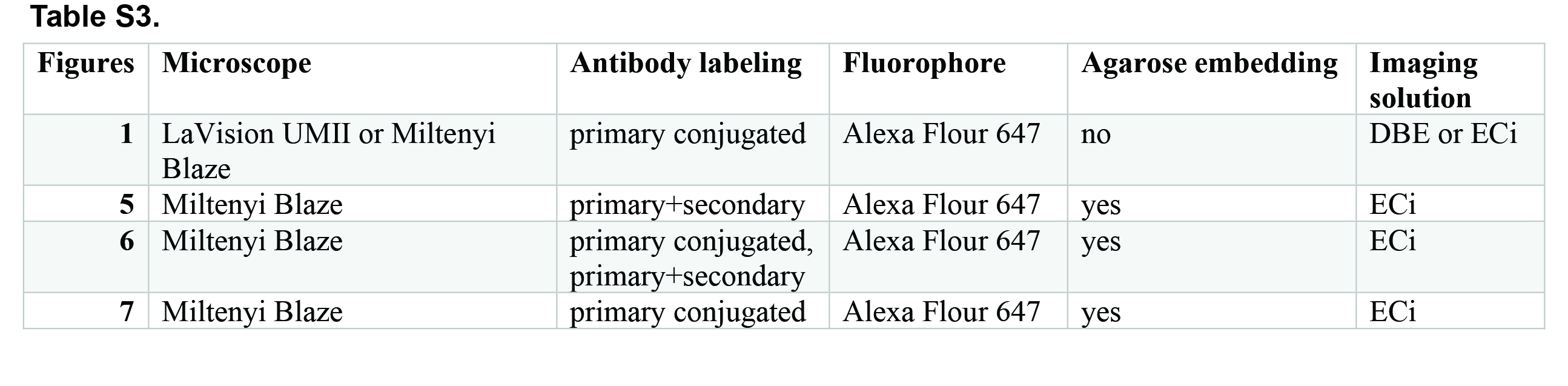
