## Supplementary material for "*Sox9* links biliary maturation to branching morphogenesis": File S1: File S1.Rmd

````
---
title: "Sholl Analysis"
output: html_document
date: "2023-12-09"
---

# Package Installation
```{r}
#Install these Packages - only need to do once
#install.packages("data.table")
#install.packages("caret")
#install.packages("ggplot2")
#install.packages("svglite")

library(data.table)
library(caret)
library(ggplot2)
library(svglite)
```

# Variable Definition
```{r}
# Sample Name
sample_name <- "Sample_2084"

# Point to data file
data <- fread(paste0("samples/",sample_name,".csv"))

# Choose the maximum polynomial fit 
max_poly_fit <- 7

# Set lower and upper percentile for linear regression on trimmed data
x_lower_percentile <- 0.05
x_upper_percentile <- 0.95

# Set random seed for consistent results (only used in cross-validation step to select best fit - shouldn't ultimately affect results)
random_seed <- 2314

# Choose whether to normalize distance
normalization <- TRUE

# Choose whether to save outputs
save_outputs <- TRUE

```

#Data Clean Up and Normalization
```{r}
# Remove rows at beginning and end where y=0
# Find the first and last indices where y is non-zero
first_non_zero <- which(data$y > 0)[1]
last_non_zero <- which(data$y > 0)[length(which(data$y > 0))]

# Filter the data to keep rows between these indices
data <- data[first_non_zero:last_non_zero, ]

# Normalize distance values (x) if user input for normalization is TRUE
if(normalization){
  data$x <- data$x / max(data$x)
  x_label <- "Normalized Distance"
} else {
  x_label <- "Distance (µm)"
}

```


#Function to find best polynomial fit 
```{r}
#Function to find best polynomial fit
findBestPolynomialFit <- function(data) {
  bestDegree <- 1
  lowestRMSE <- Inf
  bestModel <- NULL
  
  #Monte-Carlo cross-validation 
  control <- trainControl(method = "LGOCV", 
                          p = 0.8,  # Percentage of data to use for training
                          number = 20,  # Number of repetitions
                          summaryFunction = defaultSummary)
  
  for (degree in 1:max_poly_fit) {
    # Construct the formula with the current degree
    formula <- as.formula(paste("y ~ poly(x, ", degree, ", raw=TRUE)"))
    
    # Set random seed
    set.seed(random_seed)
    
    # Fit and evaluate model using Monte-Carlo cross-validation
    fit <- train(formula, 
                 data = data, 
                 method = "lm", 
                 metric = "RMSE", 
                 trControl = control)

    # Extract the RMSE for the current model
    modelRMSE <- min(fit$results$RMSE)

    # Check if this model has a lower RMSE
    if (modelRMSE < lowestRMSE) {
      lowestRMSE <- modelRMSE
      bestDegree <- degree
      bestModel <- fit
    }
  }
  cat("Best polynomial degree based on MCCV:", bestDegree, "\n")
  return(bestModel)
}

```


# Find Polynomial Fit
```{r}

# Find the best polynomial degree
bestModel <- findBestPolynomialFit(data)
bestModel_degree <- bestModel$terms[[3]][[3]]

# Calculating key metrics: Max Raw Value (Max), Critical value (Nm), Critical radius (rc)  and Mean value (Nav)

# Find the maximum raw data point
max_raw <- max(data$y)

# Create a dense grid of x-values to cover entire measured radius with step size of 0.001
x_grid <- seq(min(data$x), max(data$x), by = min(data$x)/1000)

# Predict y-values for that dense x-grid using the model (ie fitted function)
predicted_data <- data.frame(x = x_grid)
predicted_data$y <- predict(bestModel, newdata = predicted_data)

# Calculate Mean Value (Nav), Critical Value (Nm), and Critical Radius (rc) using fitted function
Nav <- mean(predicted_data[predicted_data$y > 0, ]$y) # Filter out negative y-values which may occur at extremities and skew data
max_index <- which.max(predicted_data$y)
Nm <- predicted_data$y[max_index]
rc <- predicted_data$x[max_index]

# Print the metrics
cat("Sample Name:", sample_name, "\n")
cat("Max Raw Value (Max):", max_raw, "\n")
cat("Mean Value (Nav):", Nav, "\n")
cat("Critical Value (Nm):", Nm, "\n")
cat("Critical Radius (rc):", rc, "\n")

# Predict data using original distance (x) points to graph 
predicted_data_plot <- predict(bestModel, newdata = data)

# Create the plot
p_poly  <- ggplot(data, aes(x=x, y=y)) +   # Raw measured data
  geom_point(size=1) +  # Size of measure points
  geom_line(aes(y=predicted_data_plot), color = "red", linewidth=1.5, alpha=0.7) +
  geom_segment(aes(x = rc, y = 1, xend = rc, yend = 0), 
               arrow = arrow(type = "closed", length = unit(2, "mm")), color = "grey") + # Arrowhead for rc 
  annotate("text", x = rc, y = 1.1, label = paste("rc =", round(rc, 3)), vjust = -1.0, size = 5) +  # Label for rc (rounded 1 decimal)
  
  geom_segment(aes(x = 0.01, xend = 0, y = Nav, yend = Nav), 
               arrow = arrow(type = "closed", length = unit(2, "mm")), color = "grey") + # Arrowhead for Nav
  annotate("text", x = 0.01, y = Nav, label = paste("Nav =", round(Nav, 1)), hjust = -0.1, size = 5) +  # Label for Nav (rounded 1 decimal)

  geom_hline(aes(yintercept=max_raw), linetype=2, color = "grey") +  # Dotted line indicating Maximum Raw Value
  annotate("text", x = Inf, y = max_raw, label = paste("Max =", round(max_raw, 2)), hjust = 1.1, vjust = 1.5, size = 5) +  # Label for max_raw

  geom_hline(aes(yintercept=Nm), linetype=2, color = "grey") +    # Dotted line indicating Nm (maximum fitted value)
  annotate("text", x = Inf, y = Nm, label = paste("Nm =", round(Nm, 2)), hjust = 1.1, vjust = 1.5, size = 5) +  # Label for Nm
  
  labs(x = x_label, y = "N. of intersections", title = sample_name) +
  theme_minimal() +
  theme(
    axis.line = element_line(size = 0.5),  # Bold axes lines
    axis.text.x = element_text(size = 12),
    axis.text.y = element_text(size = 12),
    axis.title=element_text(size=14),
    panel.grid.major.x = element_blank(),  # Remove grids beyond x=0
    panel.grid.minor.x = element_blank(),
    panel.grid.major.y = element_blank(),  # Remove grids beyond y=0
    panel.grid.minor.y = element_blank()
  ) +
  scale_x_continuous(expand = c(0, 0), limits = c(0, NA)) + 
  scale_y_continuous(expand = c(0, 0), limits = c(0, NA))

# Print the plot
print(p_poly)

# Define file path and save
if(save_outputs){
  
  #Save the plot
  polynomial_file_name <- paste0(sample_name, "_polynomial_regression_Sholl_plot.svg")
  
  #Save the plot
  file_path <- paste0("results/normalized/", sample_name, "/")
  polynomial_file_name <- paste0(sample_name, "_polynomial_Sholl_plot.svg")
  ggsave(path=file_path, filename=polynomial_file_name, plot = p_poly, width = 7, height = 4.5)

  }
```

# Linear Regression of Log-Transformed Data
```{r}

# Normalize and log-scale of number of intersection: log(N. intersections/area shell)
data$y_normalized <- log(data$y) - log(4 * pi * data$x^2) # Based on Log(A/B) = Log(A) - Log(B)

#Linear regression on entire range
model_full <- lm(y_normalized ~ x, data = data)

#Linear regression on middle 80 percentile
x_lower_bound <- quantile(data$x, x_lower_percentile)
x_upper_bound <- quantile(data$x, x_upper_percentile)
data_middle <- subset(data, x >= x_lower_bound & x <= x_upper_bound)
model_middle <- lm(y_normalized ~ x, data = data_middle)

# Generate plot
p_log <- ggplot(data, aes(x = x, y = y_normalized)) +
  #geom_rect(aes(xmin = x_lower_bound, xmax = x_upper_bound, ymin = -Inf, ymax = Inf), fill = "grey95", alpha = 0.5) +
  geom_point(size = 1) +  # Plot the normalized data points
  geom_smooth(method = "lm", formula = y ~ x, color = "blue", data = data) +
  #geom_smooth(method = "lm", formula = y ~ x, color = "red", data = data_middle) +
  labs(x = x_label, y = expression("N. of intersections /" ~µm^2 ~ " (log)"), title = sample_name) +
  theme_minimal() +
  theme(
    axis.line = element_line(size = 0.5),  # Bold axes lines
    axis.text.x = element_text(size = 12),
    axis.text.y = element_text(size = 12),
    axis.title=element_text(size = 14),
    panel.grid.major.x = element_blank(),  # Remove grids 
    panel.grid.minor.x = element_blank(),
    panel.grid.major.y = element_blank(),  # Remove grids 
    panel.grid.minor.y = element_blank()
  )

# Print the plot
print(p_log)

# Output the Sholl coefficients
k_full <- -1 * coef(model_full)["x"] # Sholl coefficient for entire data
k_middle <- -1 * coef(model_middle)["x"] # Sholl coefficient for middle 80% of data

# Extract the R-squared values
summary_full <- summary(model_full)
R_squared_full <- summary_full$r.squared

summary_middle <- summary(model_middle)
R_squared_middle <- summary_middle$r.squared

# Print the metrics
lower_percentile <- x_lower_percentile*100
upper_percentile <- x_upper_percentile*100
percentile_range <- paste0(lower_percentile," - ",upper_percentile)
cat("Sample Name:", sample_name, "\n")
cat("R squared, full data", R_squared_full, "\n")
cat("Sholl coefficient, full data:", k_full, "\n")
cat(paste0("R squared, ",lower_percentile,"th-",upper_percentile,"th percentile: ", R_squared_middle), "\n")
cat(paste0("Sholl coefficient, ",lower_percentile,"th-",upper_percentile,"th percentile: ", k_middle), "\n")


# Define file path and save
if(save_outputs){

  #Save the plot
  file_path <- paste0("results/normalized/", sample_name, "/")
  log_file_name <- paste0(sample_name, "_log_Sholl_plot.svg")
  ggsave(path=file_path, filename= log_file_name,plot = p_log, width = 7, height = 4.5)
  
  }
```

# Create output data table
```{r}
# Create a data frame
output_table <- data.frame(
  Variable = c("Sample Name", "Best Polynomial Model Degree", "Normalized",
               "Max Raw Value", "Mean Value (Nav)", "Critical Value (Nm)", 
               "Critical Radius (rc)", "R Squared (Full)", "Sholl Coefficient (Full)", 
               "Percentile Range", "R Squared (Percentile Range)", "Sholl Coefficient (Percentile Range)"),
  Value = c(sample_name, bestModel_degree, normalization, 
            max_raw, Nav, Nm, rc, 
            R_squared_full, k_full, 
            percentile_range, R_squared_middle, k_middle)
)

# Define file path and save
if(save_outputs){

  output_file_path <- paste0("results/normalized/", sample_name, "/",sample_name, "_Sholl_output_variables.csv")
  write.csv(file = output_file_path, output_table, row.names = FALSE)

  }
```
````
